# The functional form of value normalization in human reinforcement learning

**DOI:** 10.1101/2022.07.14.500032

**Authors:** Sophie Bavard, Stefano Palminteri

## Abstract

Reinforcement learning research in humans and other species indicates that rewards are represented in a context-dependent manner. More specifically, reward representations seem to be normalized as a function of the value of the alternative options. The dominant view postulates that value context-dependence is achieved via a divisive normalization rule, inspired by perceptual decision-making research. However, behavioral and neural evidence points to another plausible mechanism: range normalization. Critically, previous experimental designs were ill-suited to disentangle the divisive and the range normalization accounts, which generate similar behavioral predictions in many circumstances. To address this question, we designed a new learning task where we manipulated, across learning contexts, the number of options and the value ranges. Behavioral and computational analyses falsify the divisive normalization account and rather provide support for the range normalization rule. Together, these results shed new light on the computational mechanisms underlying context-dependence in learning and decision-making.

## Introduction

The process of attributing economic values to behavioral options is highly contextdependent: the representation of an option’s utility does not solely depend on its objective value, but is strongly influenced by its surrounding (i.e., other options simultaneously or recently presented). This is true in an extremely wide range of experimental paradigms, ranging from decision among lotteries to reinforcement learning problems [1–5]. This is also true for a wide variety of species, including mammals [6,7], birds [8] and insects [9, 10]. The pervasiveness of this effect across tasks and species suggests that context-dependence may reflect the way neuron-based decision systems address a fundamental computational trade-off between behavioral performance and neural constraints [11–14].

Indeed, it has been showed that context-dependence often takes the form of a normalization process where option values are rescaled as a function of the other available options, which has the beneficial consequence of adapting the response to the distribution of the outcomes [14]. the idea that neural codes and internal representations are structured to carry as much as information per action is the cornerstone of the efficient coding hypothesis, demonstrated both at the behavioral and neural levels, in perceptual decision-making [15].

Probably due to its popularity in perception neuroscience, the dominant view regarding the computational implementation of value normalization in economic decisions postulates that is follows a divisive rule, according to which the subjective value of an option is rescaled as a function of the sum of the value of all available options [16–19]. In addition, to be validated in the perceptual domain, the divisive normalization rule also presents the appeal of being reminiscent of Herrnstein’s matching law for behavioral allocation [20]. However, while dominant, the divisive normalization account of value normalization, is not consensual [12, 13, 21–23]. Indeed, range normalization represents a plausible alternative account of value normalization and is made plausible by both behavioral and neural observations [24, 25]. According to the range normalization rule, option values are rescaled as a function of the maximum and the minimum values presented in a context, irrespective of the number of options (or choice set size) [7, 24, 26]. Answering this question bears important consequences for neuroscience, because understanding the scaling between objective and subjective outcomes is paramount to investigate the neural codes of economic values and understand the neural mechanisms of decision-making [27, 28]. Yet, the experimental paradigms used so far were ill-suited to distinguish between two accounts of value normalization in learning and decision-making [3]. To address this issue, we designed a new reinforcement learning protocol where, by simultaneously manipulating outcome ranges and the number of options, we made the divisive and the range normalization predictions qualitatively diverge in many respects [29, 30]. We opted for a reinforcement learning paradigm because it has a greater potential for translational and cross-species research [31]. In a total of six experiments (N=50 each) we deployed several variations of this new behavioral protocol where we controlled for several factors. The behavioral, model fitting and simulations results convergently rejected divisive normalization as a satisfactory explanation of the results in favor of the range normalization account. Results also suggested that the range normalization account should be further improved by non-linear weighting process. To check the robustness of our results across different elicitation methods and representational systems, we also assessed option values using explicit ratings. Values inferred from explicit, declarative, ratings were remarkably consistent with those inferred from more traditional, choice-based, methods.

## Results

### Computational hypotheses and *ex-ante* model simulations

The goal of this study was to characterize the functional form of outcome (or reward) normalization in human reinforcement learning. More specifically, we aimed at arbitrating between two equally plausible hypotheses: range normalization and divisive normalization. Both hypotheses assume that after reception of a given objective reward *R*, the learner forms an internal, subjective, representation of it, *R*_NORM_, which is influenced by other contextually relevant rewards. Crucially, the two models differ in how *R*_NORM_ is calculated. According to the range normalization hypothesis, the subjective normalized reward *R*_NORM_ is defined as the position of the objective reward R within its contextual range

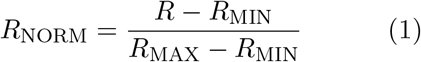

where *R*_MAX_ and *R*_MIN_ are the endpoints of the contextually relevant distribution and together form the range (*R*_MAX_−*R*_MIN_). On the other side, the divisive-normalization hypothesis, in its simplest form, postulates that the subjective normalized reward *R*_NORM_ is calculated by dividing the objective reward by the sum of all the other contextually rewards:

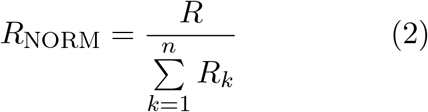

where *n* is the number of contextually relevant stimuli. These hypotheses concerning value normalization are then easily plugged into the reinforcement learning framework, simply by assuming that the value of an option is updated by minimizing a prediction error calculated on the basis of the subjective reward. Although these normalization functions are mathematically distinct, they make identical (or very similar) behavioral predictions in many of the experimental protocols designed to investigate context-dependent reinforcement learning so far [3–5, 26, 32]. For the present study we designed reinforcement learning tasks perfectly suited to adjudicate these computational hypotheses. The key idea of our behavioral protocol is to orthogonally manipulate across different learning contexts the amplitude of the range of the possible outcomes and the number of options (often referred to as choice size set; **Figure 1A**; see also **Figure S1A** for an alternative version). The first factor (the amplitude of the range of the possible outcomes) is key to differentiate an unbiased model (where *R*_NORM_ = *R*) from both our normalization models. The second factor (number of options) is key to differentiate the range normalization from divisive normalization. The reason for this can easily be inferred from **Equations 1 and 2**, because adding more option outcomes has a significant impact on the subjective reward *R*_NORM_ only following the divisive-normalization rule ^1^. To quantitatively substantiate these predictions, we run model simulations using three models. We compared a standard model with unbiased subjective values (UNBIASED), and two normalization models using either the divisive or the range normalization rules (referred to as DIVISIVE and RANGE, respectively). First, we simulated a learning phase, where each learning context in our factorial design was presented 45 times. After each trial the simulated agent was informed about the outcomes that were drawn from normal distributions (**Figure 1B**). To avoid sampling issues and ambiguity concerning the definition of the relevant normalization variables, the simulated agents were provided information about the outcomes of all options (‘complete’ feedback [34, 35]). After the learning phase, the simulated agents went through a transfer phase, where they made decisions among all possible binary combinations of the options (without additional feedback being provided). Similarly constructed experiments, coupling a learning to a transfer phase, have been proven key to demonstrate contextual effects in previous studies [3, 4, 8, 26, 32, 36].

**Figure 1:**
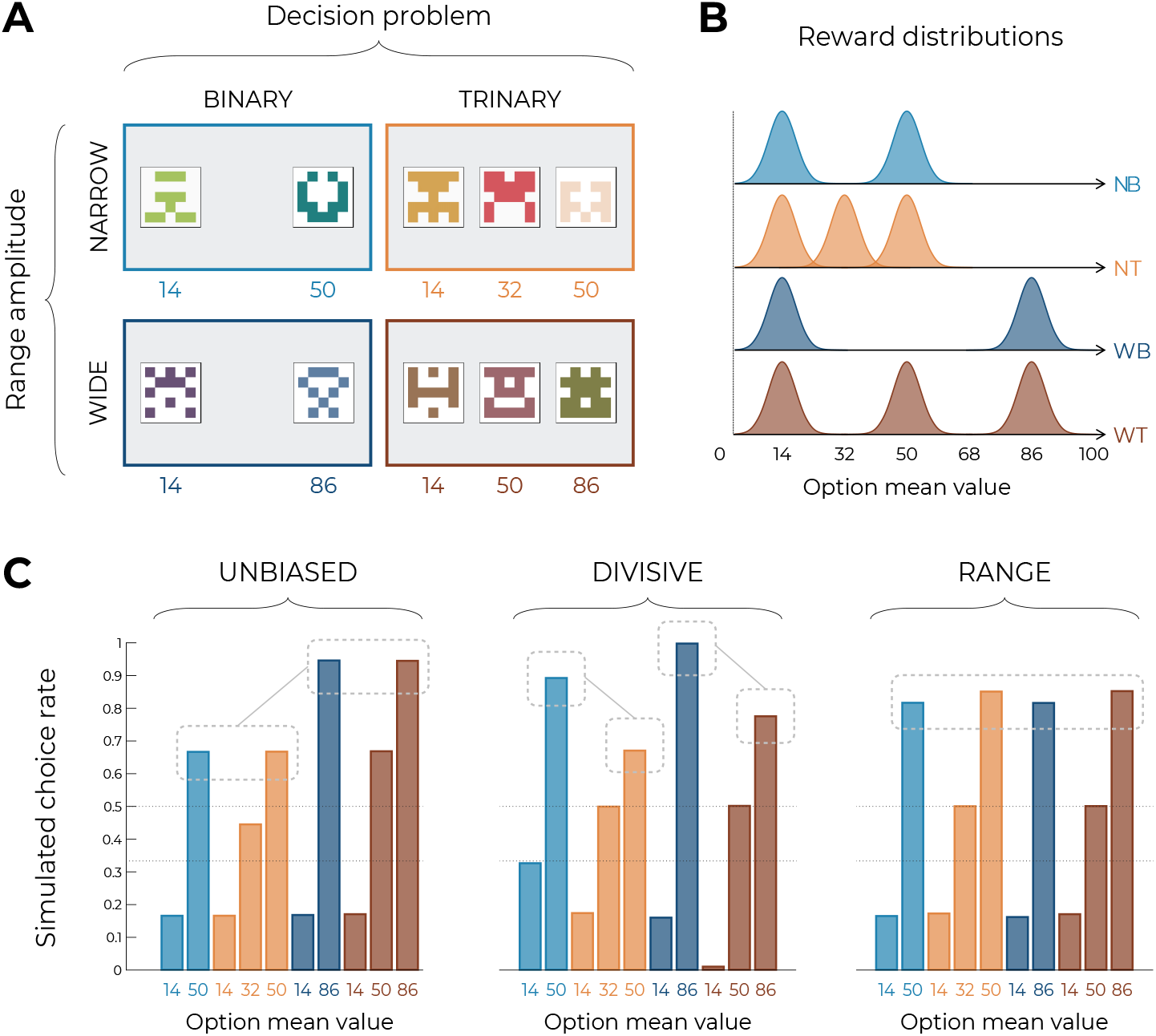
Experimental design and model predictions. **(A)** Choice contexts in the learning phase. Participants were presented with four choice contexts varying in the amplitude of the outcomes’ range (narrow or wide) and the number of options (binary or trinary decisions). **(B)** Means of each reward distribution. After each decision, the outcome of each option was displayed on the screen. Each outcome was drawn from a normal distribution with variance *σ*^2^ = 4. NB: narrow binary, NT: narrow trinary, WB: wide binary, WT: wide trinary. **(C)** Model predictions of the transfer phase choice rates for the UNBIASED (left), DIVISIVE (middle) and RANGE (right) models. Dashed lines represent the key prediction for each model.

When analyzing model simulations, we focus on choices patterns in the transfer phase (of accuracy during the learning phase is weakly diagnostic because all models predict above chance accuracy, whose level depends the choice stochasticity). **Figure 1C** plots the average simulated choice rate in the transfer phase. It is calculated across all the comparisons involving a given option. Unsurprisingly, within each learning contexts, in all models the choice rates are higher for high value options compared to lower value options. However, model simulations show that the models produce choice patterns that differ in many key aspects. Let’s start considering the UNBIASED model as a bench mark (**Figure 1C**, left). Since it encodes outcomes in an unbiased manner, it predicts higher choice rate for the high value option in the ‘wide’ contexts (WB_86_ and WT_86_) compared to high value options in the ‘narrow’ contexts (NB_50_ and NT_50_). On the other side, the UNBIASED model predicts that choice rate in the transfer phase is not affect by whether or not the option belonged to a binary or a trinary learning context. Moving to the DIVISIVE model, we note that the difference between the choice rates of high value options of the ‘wide’ contexts (WB_86_ and WT_86_) compared those of the ‘narrow’ contexts (NB_50_ and NT_50_) is now much smaller, due to the normalization process (**Figure 1C**; middle). However, the DIVISIVE model also predicts that the choice rate is strongly affected by whether or not the option belonged to a binary or a trinary learning context. For instance, WB_86_ and NB_50_ present a much higher choice rate compared to WT_86_ and NT_50_, respectively, despite their objective expected value being the same. This is an easily identifiable and direct consequence of the denominator of the divisive formulation rule increasing as a function of the number of options (see **Equation 2**). Concerning the RANGE model (**Figure 1C**; right), it predicts choice rates being similar across all high value options, regardless of their objective values (because of the normalization) and whether or not the option belonged to a trinary or binary context (because of the range normalization rule). Finally, the choice rates of the low value (14) options also discriminate the DIVISIVE model, where it is strongly modulated by the task factors, from the other two models, where all low value options present the same choice rate. To conclude, model simulations confirm that our design, involving a factorial learning phase and a transfer phase, is well suited to disentangle our three a priori models because they predict qualitatively differentiable patterns of choices (see also **Figure S1C** for similar conclusions based on an alternative task design) [30, 37].

### Behavioral results

The above-described behavioral protocol was administered to N=50 participants recruited online which played for real monetary incentives as previously described [26]. We first tested whether the correct choice rate (i.e., the probability of choosing the option with the highest expected value) was overall above chance level during the learning phase, to check that the participants engaged in the task. Indeed, correct response rate was significantly higher than chance level (0.5 and 0.33 in the binary and trinary learning contexts, respectively) in all conditions (least significant: *t*(49) = 18.93, *p <* .0001, *d* = 2.68; on average: *t*(49) = 24.01, *p <* .0001, *d* = 3.96; **Figure 2A**). We further checked whether the task factors affected performance in the learning phase and found a significant effect of the decision problem (the correct choice rate being higher in the binary compared to the trinary contexts: *F* (1, 49) = 9.26, *p* = .0038, *η*^2^ = 0.16), but no effect of range amplitude (wide versus narrow; *F* (1, 49) = 0.52, *p* = .48, *η*^2^ = 0.01) or interaction (*F* (1, 49) = 2.23, *p* = .14, *η*^2^ = 0.04). We next turned to the results from the transfer phase. Following the analytical strategy used in previous studies, we first checked that the correct choice rate in the transfer was significantly higher than chance (*t*(49) = 9.10, *p <* .0001, *d* = 1.29), thus providing positive evidence of value retrieval and generalization [4,26,36]. We analyzed the choice rate per symbol, which is the average frequency with which a given symbol is chosen in the transfer phase, and can therefore be taken as a measure of the subjective preference for a given option [4, 32]. We focus on key comparisons that crucially discriminate between competing models of normalization. First, and contrary to what was predicted by the DIVISIVE model, the choice rate for the highest value options of in the trinary contexts (NT_50_ and WT_86_) was not lower compared to that of the binary ones (NB_50_ and WB_86_). Indeed, if anything, their choice rate was higher (NT_86_ vs. NB_86_: *t*(49) = 1.66, *p* = .10, *d* = 0.29; WT_86_ vs. WB_86_: *t*(49) = 2.80, *p* = .0072, *d* = 0.53). Similarly, the choice rate of the lowest value options was not affected by the by their belonging to a binary or trinary context in the direction predicted by the DIVISIVE model. For instance, if anything the choice rate for the WT_14_ was numerically (but not significantly) lower compared that of NB_14_ (*t*(49) = −0.11, *p* = .92, *d* = −0.022). Concerning other features of the transfer phase performance, some comparisons were superficially consistent with the UNBIASED model, such as the fact that highest value options in the narrow contexts (NB_50_ and NT_50_) displayed a lower choice rate compared to the highest value options of the wide contexts (NB_86_ and NT_86_; *t*(49) = −4.19, *p* = .00011, *d* = −0.72), even if the size of the difference appeared to be much smaller to that expected from ex ante model simulations (**Figure 1C**, right). Other features were clearly more consistent with the RANGE model. For instance, the fact that the mid value option in the wide trinary context WT_50_ displayed a significantly lower choice rate compared to the highest value options in the narrow contexts (NT_50_ and NB_50_) was not predicted by UNBIASED. Finally, the mid value options (NT_32_ and WT_50_) displayed a choice rate very close to that of the corresponding lowest value options (NT_14_ and WT_14_): this feature is clearly in contrast with the DIVISIVE and UNBIASED models (which predict their choice rate closer to that of the corresponding highest value options: NT_50_ and WT_86_), but not perfectly captured either by the RANGE model (which predicts their choice rate perfectly in between those of high and low value options). To rule out that this effect was not due to a lack of attention for the low and mid value options, we designed two additional experiments where we added forced choice trials to focus the participants’ attention on all possible options (**Table 1**) [38]. In one experiment (N=50), forced choice trials were followed by partial feedback (**Figure 2B**), in another experiment (N=50) forced choice trials were followed by partial feedback (**Figure 2C**). Focusing participants’ attention to all possible outcomes by forcing their choice did not significantly affect the behavioral performance neither in the learning phase (*F* (2, 147) = 2.75, *p* = .067, *η*^2^ = 0.04, Levene’s test *F* (2, 147) = 2.43, *p* = .092) nor in the transfer phase (*F* (2, 147) = 0.64, *p* = 53, *η*^2^ = 0.00, Levene’s test *F* (2, 147) = 0.64, *p* = .53). This suggests that the choice rates of the mid options reflect their underlying valuation (rather than lack of information). Given the absence of detectable differences across experiments, in the model-based analyses that follow, we pooled the three experiments together. To sum up, behavioral results, specifically in the transfer phase, are in contrast with the predictions of the DIVISIVE model and are rather consistent with the range normalization process proposed by the RANGE model. Behavioral results in three experiments (N=50 each) featuring a slightly different design, converge to the same conclusion (see **Supplementary materials**). In the following section, we substantiate this claim by formal model comparison and ex post model simulations analysis.

**Table 1:** Experimental design. Each version of each experiment was composed of four different learning contexts. Results of experiment 1 are presented in the main text; results of experiment 2 are presented in the **Supplementary materials**.

|  | N | Learning contexts |  |  |  |  |  | N forced trials (feedback) |
| --- | --- | --- | --- | --- | --- | --- | --- | --- |
|  |  | [14,50] | [14,32,50] | [50,86] | [50,68,86] | [14,86] | [14,50,86] |  |
| Experiment 1a | 50 | X | X |  |  | X | X | 0 |
| Experiment 1b | 50 | X | X |  |  | X | X | 0 |
| Experiment 1c | 50 | X | X |  |  | X | X | 50 (complete) |
| Experiment 2a | 50 |  |  | X | X | X | X | 50 (complete) |
| Experiment 2b | 50 |  |  | X | X | X | X | 50 (partial) |
| Experiment 2c | 50 |  |  | X | X | X | X | 50 (partial) |

**Figure 2:**
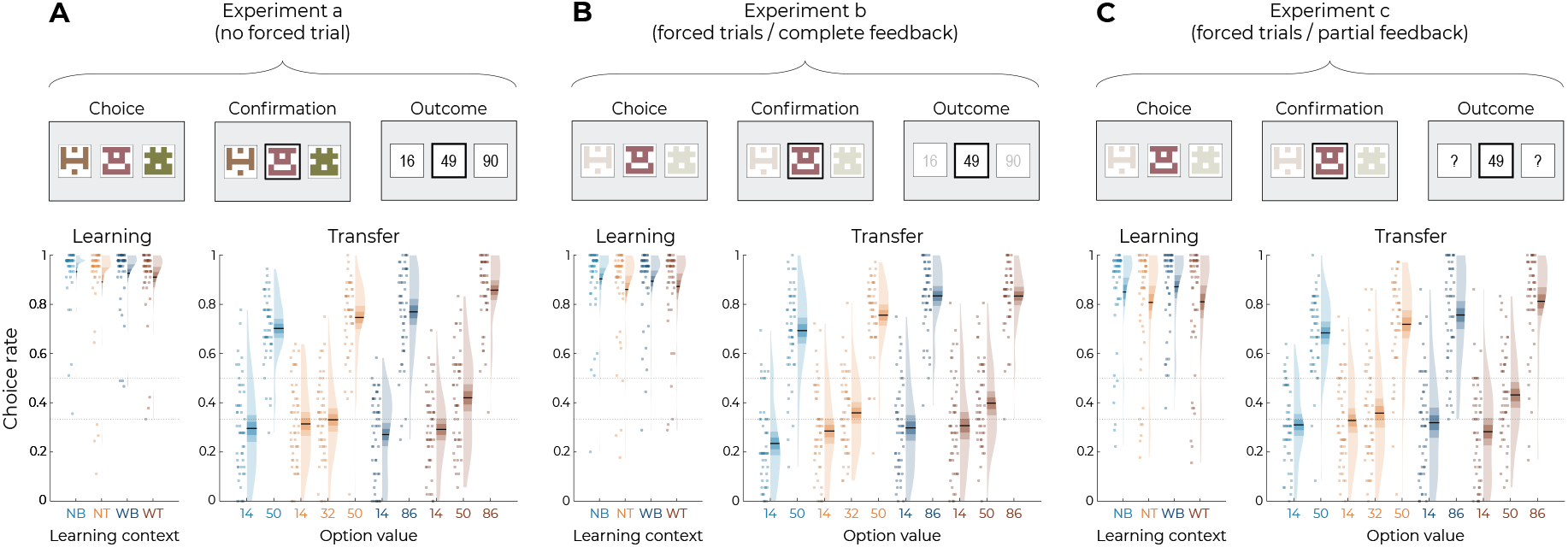
Behavioral results. Top: successive screens of a typical trial for the three versions of the main experiment: without forced trials **(A)**, with forced trials and complete feedback information **(B)** and with forced trials and partial feedback information **(C)**. Bottom: correct choice rate in the learning phase as a function of the choice context (left panels), and choice rate per option in the transfer phase (right panels) for the three versions of the main experiment: without forced trials **(A)**, with forced trials and complete feedback information **(B)** and with forced trials and partial feedback information **(C)**. In all panels, points indicate individual average, shaded areas indicate probability density function, 95% confidence interval, and SEM. NB: narrow binary, NT: narrow trinary, WB: wide binary, WT: wide trinary.

### Model comparison and ex post model simulations

Behavioral analyses of transfer phase choices suggest that learning and valuation are more consistent with the predictions of the RANGE model, compared to those of the UNBIASED or the DIVISIVE model. To quantitatively substantiated this claim, we formally compared the quality of fit of the three models using and out-of-sample log-likelihood [39].

Specifically, we first optimized the models’ free parameters (learning rates and choice inverse temperature) in order to maximize the log-likelihood of observing the learning phase choices, given the model and the parameters. We then used these parameters to generate the log-likelihood of observing the choices in the transfer phase, which were not included in the original model fitting. The RANGE model displayed a much higher mean and median log-likelihood (which indicated better fit) compared to both the DIVISIVE and the UNBIASED models (oosLL_RAN_ vs oosLL_DIV_: *t*(149) = 10.10, *p <* .0001, *d* = 0.41; oosLL_RAN_ vs oosLL_UNB_: *t*(149) = 8.34, *p <* .0001, *d* = 0.82; **Table 2**). Subsequently, we simulated transfer choice phase using the best fitting, i.e., empirical, parameter values (**Figure 3**). The results of this ex post simulations confirm what inferred from the ex-ante simulations and indicate that RANGE model predicted results much closer to the observed ones in respect of many key comparisons. All these results were replicated in three additional experiments feature slightly different design, where the DIVISIVE model displayed a higher mean log-likelihood compared to the UNBIASED model, indicating no robust improvement of the quality of fit (see **Table 2** and **Supplementary Materials**). Despite the superiority of the RANGE model in terms of both predictive (out-of-sample log-likelihood) and generative (simulation) performance [39] compared to UNBIASED and DIVISIVE one, it still failed to perfectly captured transfer phase preference, specifically concerning the mid value options. In the subsequent section we propose how the RANGE model could be further improved to obviate this issue.

**Table 2:**
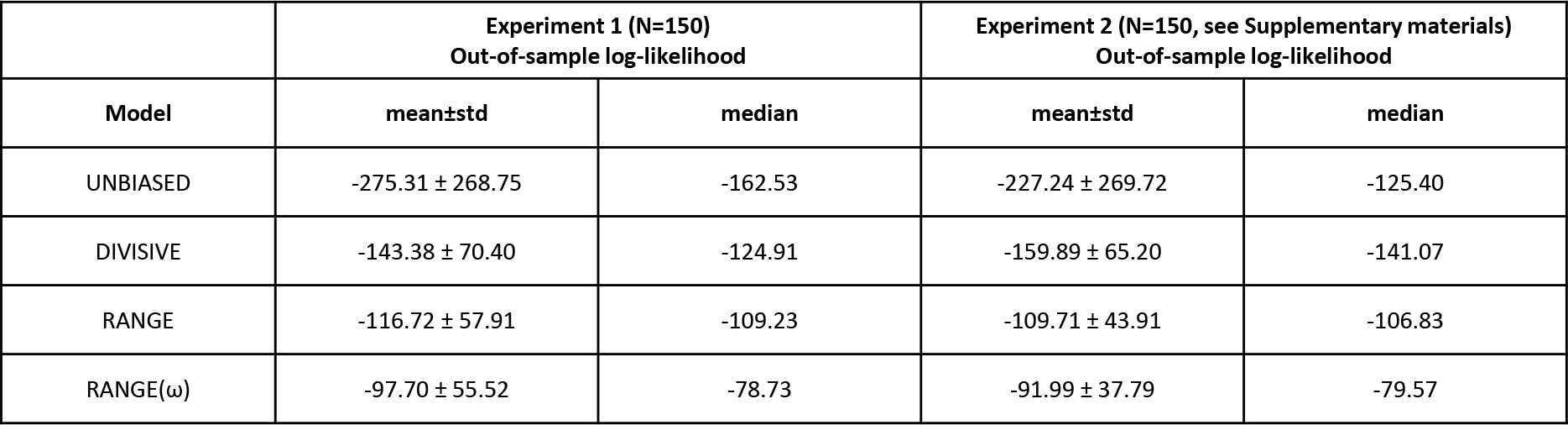
Quantitative model comparison. Values reported here represent mean *±* std and median of out-of-sample log-likelihood for each model.

**Figure 3:**
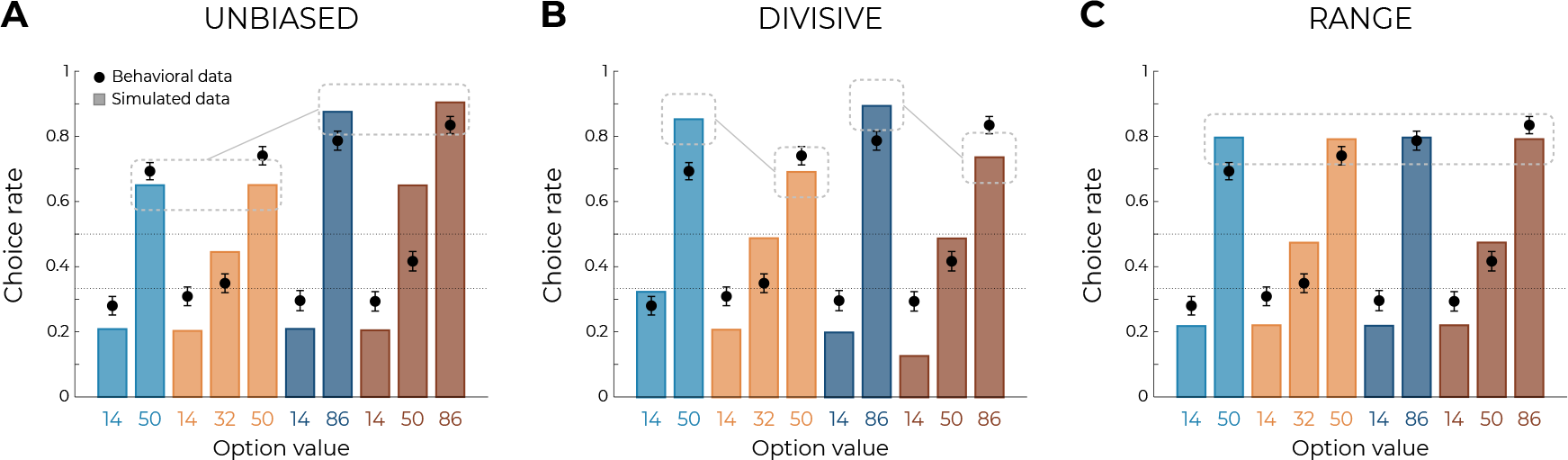
Qualitative model comparison. Behavioral data (black dots) superimposed on simulated data (colored bars) for the UNBIASED **(A)**, DIVISIVE **(B)** and RANGE **(C)** models. Simulated data in the transfer phase were obtained with the best-fitting parameters, optimized on all four contexts of the learning phase. Dashed lines represent the key prediction for each model.

### Improving the RANGE model

Although model comparison and model simulation both unambiguously favored the RANGE model over the UNBIASED and DIVISIVE models, the RANGE model is not perfect at predicting participants’ choices in the transfer phase (**Figure 3C**). As mentioned previously, the mid value options in trinary contexts (NT_32_ and WT_50_) displayed a choice rate closer to that of the corresponding lowest value options (NT_14_ and WT_14_): a feature which was not captured by the RANGE model (it predicts their choice rate to be exactly halfway of those of low and high value options (NT_86_ and WT_86_). In addition, the choice rate of lowest value options of all contexts (NB_14_, NT_14_, WB_14_ and WT_14_) was underestimated by the RANGE model. These observations are *prima facie* compatible with the idea that outcomes are not processed linearly [40, 41]. To formally test this hypothesis with the goal of improving the RANGE model, we augmented it with a free parameter *ω* which applies a non-linear transformation to the normalized outcome. More specifically, in this modified RANGE model (RANGE^*ω*^), the normalized outcome is power-transformed by the *ω* parameter (0 *< ω <* +∞), as follows:

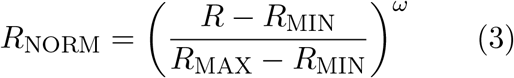

Crucially, for *ω* = 1, the RANGE^*ω*^ reduced to the RANGE model; for *ω <* 1, the RANGE^*ω*^ model induces a concave deformation of the normalized outcome; for *ω >* 1, the RANGE^*ω*^ model induces a convex deformation of the normalized outcome. On average, the power parameter *ω* was greater than 1 (mean±std: 2.97 ± 1.36, *t*(149) = 17.81, *p <* .0001, *d* = 1.45), suggesting that participants value the mid value options less than the midpoint between lowest and highest options (i.e., closer to the lowest option, **Figure 4A**). Quantitative model comparison favored the RANGE^*ω*^ model over all other models, including the RANGE model (**Table 2**) (oosLL_RAN_ vs oosLL_RAN_*ω* : *t*(149) = −6.98, *p <* .0001, *d* = −0.57; **Table 2**). Moreover, the inspection of model simulations confirmed that the RANGE^*ω*^ model perfectly captures participants’ behavior in the transfer phase. More specifically, the mid value options (NT_32_ and WT_50_) and the lowest value options (NB_14_, NT_14_, WB_14_ and WT_14_) were better estimated in all contexts (**Figure 4B**). To conclude, the addition of a power parameter allowed our model to match participants’ behavior almost perfectly.

**Figure 4:**
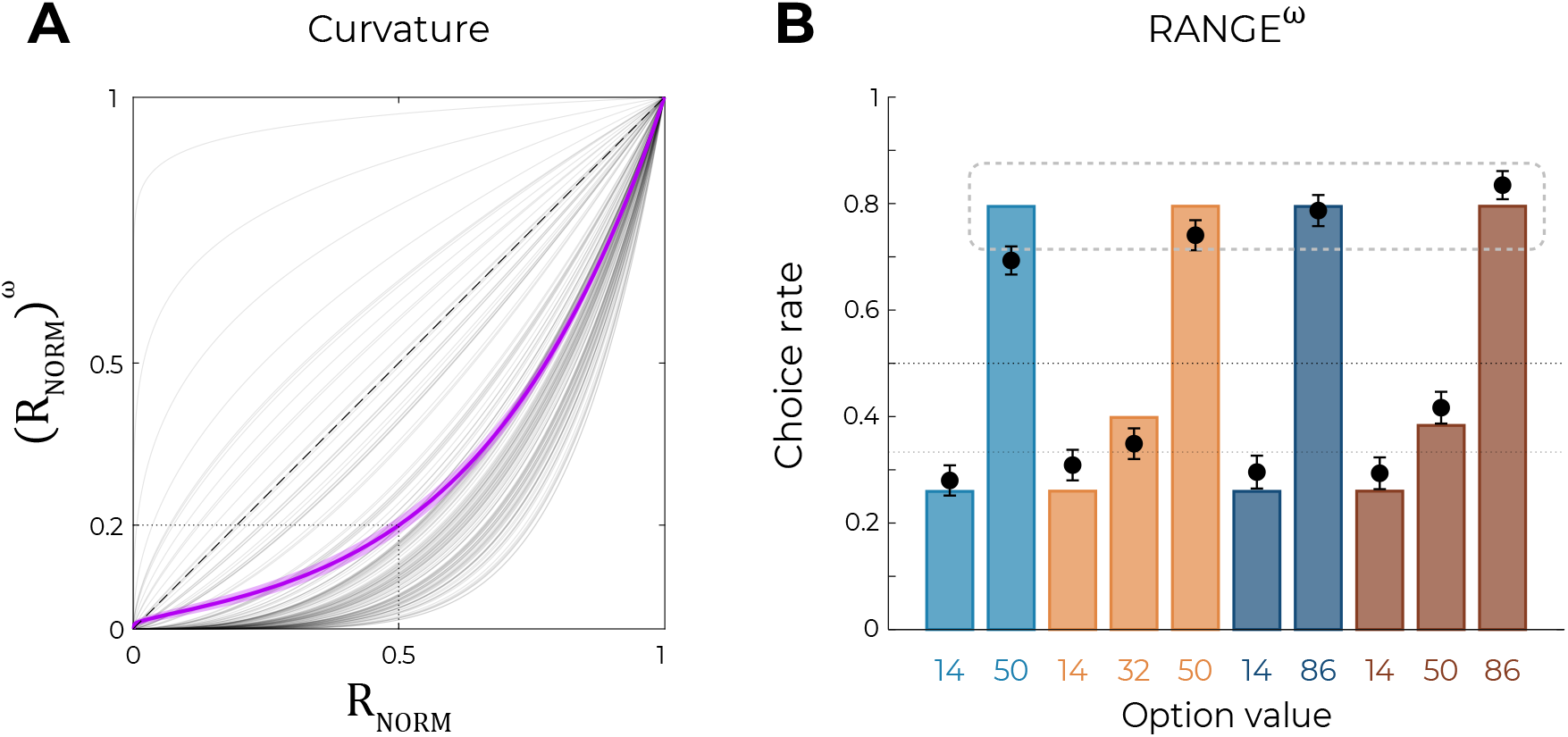
Predictions of the non-linear RANGE model. **(A)** Curvature function of the normalized reward per participant. Each grey line was simulated with the best-fitting power parameter *ω* for each participant. Dashed line represents the identity function (*ω* = 1), purple line represents the average curvature over participant, shaded area represents SEM. **(B)** Behavioral data (black dots) superimposed on simulated data (colored bars) for the RANGE^*ω*^ model. Simulated data in the transfer phase were obtained with the best-fitting parameters, optimized on all four contexts of the learning phase. Dashed lines represent the key prediction for the model.

### Explicit assessment of option values

In addition to the transfer phase, participants performed another elicitation assessment, where they were asked to explicitly rate the average value of each option using a slider ranging from 0 to 100. This explicit elicitation phase allowed us to have a complementary estimation of participants’ subjective valuations of each option, to compare them with the choice-based transfer phase. The subjective values elicited through explicit ratings were consistent with those elicited though binary choices in many key aspects (**Figure 5A**). Indeed, against what predicted by DIVISIVE principle, option subjective values did not depend on the number of options in each context, but rather on their ordinal value within the context (minimum, mid, maximum). This pattern is even clearer when looking at the difference between reported subjective values and the objective values of each option (**Figure 5B**). Crucially, the subjective value of the options with an objective value of 50 (NB_50_, NT_50_, and WT_50_) is completely determined by its position in the range of its context, and not by the total sum of the options in this context. Finally, to compare elicitation methods, we simulated transfer phase choices based on the explicit elicitation ratings. More specifically, for each participant and comparison, we generated choices using an argmax selection rule on the subjective values they explicitly reported for each option (see Equation 11). We found the pattern simulated using explicit ratings to perfectly match the actual choice rates of the transfer phase (*t*(149) = 1.51, *p* = .13, *d* = 0.12, **Figure 5C**), suggesting that both elicitation methods tap into the same valuation system. Similar results and conclusions could be drawn from Experiment 2 (see **Supplementary Materials, Figure S2**).

**Figure 5:**
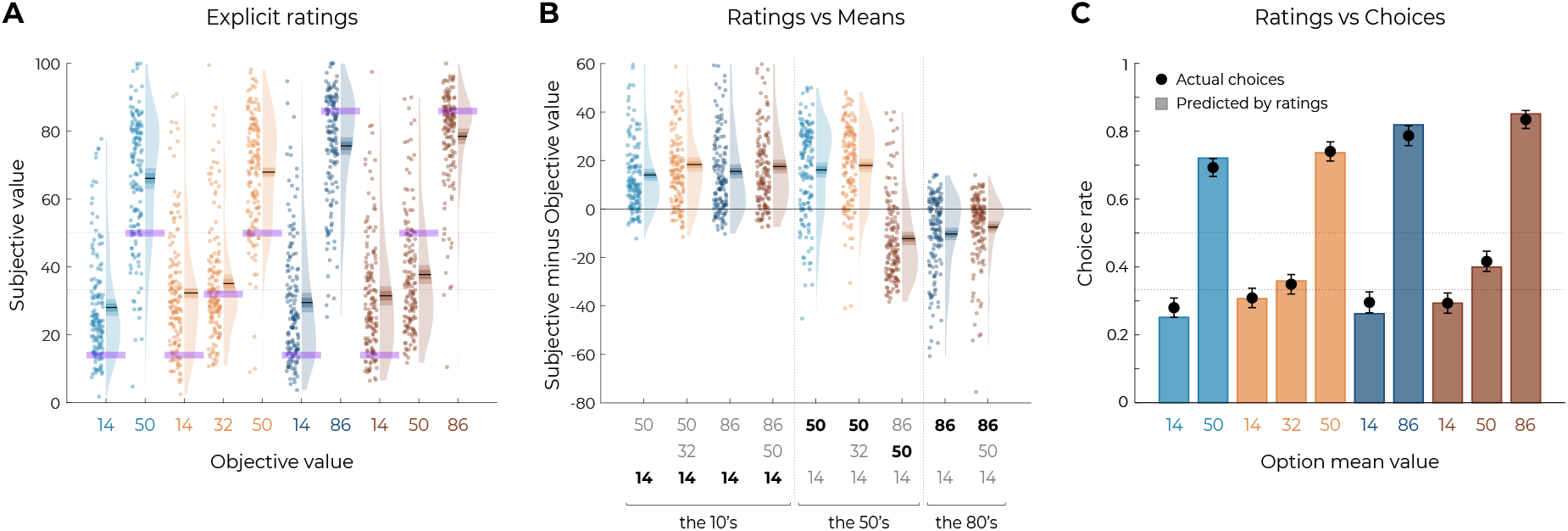
Results from the elicitation phase. **(A)** Reported subjective values in the elicitation phase for each option. Points indicate individual average, shaded areas indicate probability density function, 95% confidence interval, and SEM. Purple areas indicate the actual objective value for each option. **(B)** Difference between reported subjective value and actual objective value for each option, arrange in ascending order. The legend of the x-axis represents the values of the options of the context in which each option was learned (actual option value shown in bold black). Points indicate individual average, shaded areas indicate probability density function, 95% confidence interval, and SEM. **(C)** Behavioral choice-based data (black dots) superimposed on simulated choice-based data (colored bars). Simulated data were obtained with an argmax rule assuming that subjects were making transfer test decision based on the explicit subjective ratings of each option.

## Discussion

In the present paper we sought to challenge the current dominant view of how value-related signals are encoded for economic decision-making. More precisely we designed behavioral paradigms perfectly tailored to discriminate between unbiased (or “absolute”) representations, from context-dependent or normalized representations following different, antagonist, views. To do so, we deployed a series of six behavioral reinforcement learning experiments consisting in an initial learning phase (where participants learned to associate options to rewards) and a transfer – or generalization phase allowing us to infer the subjective learned value of each option [42].

By systematically manipulating outcome ranges, we were able to confirm that behavioral data was inconsistent with the idea that humans learn and represent values in an unbiased manner. Indeed, subjective values were similar for options presented in the decision contexts with narrow or wide decision ranges, despite their objective values being very different. This result was quantitatively backed-up by both model simulations and out-of-sample likelihood analyses that suggested the unbiased model being worst on average than any other normalization model. Thus, our findings significantly add up to the now overwhelming body of evidence indicating that value representations are highly context-dependent even in the reinforcement learning scenario, both in human [3–5, 26, 32, 36, 43] and non-human animals [6–8, 10, 44, 45].

On the other side, by contrasting binary to trinary options decision problems, we were able to provide incontrovertible evidence of the popular idea that context-dependence follows the functional form of divisive normalization, also in the context of value-based decision-making [14, 18, 33]. Our demonstration relied on the straightforward and well accepted idea that virtually any instantiation of the divisive normalization model would predict a strong choice set size effect: all things being equal, the subjective value of an option in a trinary decision context should be lower compared that of a similar option belonging to a binary context. We find no evidence for such an effect. In fact, if anything, the subjective values of options belonging to trinary decision contexts were numerically higher compared to that of the binary decision contexts. Beyond the quantitative analysis of behavioral performance, model fitting and simulations analysis also revealed that the divisive normalization model dramatically failed to correctly reproduce the behavioral pattern [30]. Crucially, this was also true for fully parametrized versions of the divisive normalization model (presented in the **Supplementary Materials** [19]).

The behavioral results were rather consistent with an alternative rule, range normalization, according to which subjective value signals are normalized as a function of the maximum and minimum values (regardless of the number of options). This normalization rule is reminiscent of the range principle proposed by Parducci to explain perceptual (and later also affective) judgements [46]. As opposite to divisive normalization, range normalization predicts that contextually high and low value options are assigned the same subjective value, regardless of their absolute value and the number of options in a given choice set. This behavior hallmark of range normalization was replicated in all experiments. Of note, although even though the quality of the fit of the range normalization model was significantly better compared that of the divisive normalization model, in terms of both predictive accuracy (out-of-sample likelihood) and behavioral signatures, it manifestly failed to properly capture the subjective values attributed to the mid-value options in trinary choice contexts. This pattern was not affected by introducing forced choice trials to focus attention of the participant to mid-option outcomes (for at least five trials) [38]. To further improve the range normalization model fit, we endowed it with a non-linear weighting of normalized subjective values. Non-linear weighting parameters, although they do not provide mechanistic accounts, are often introduced in models of decision-making to satisfactorily account for utility and probability distortions [34, 47]. Introducing this weighting parameter improved the model fit, qualitatively and quantitatively in a substantial manner. Importantly, introducing the same parameter to the divisive model did not improve its performance significantly (see Supplementary Materials). The weighted range normalization model improved the fit assuming that the mid and low value options are subjectively perceived as much closer than they are in reality. We believe that this may derive from attentional mechanisms that bias evidence accumulation as a function of outcomes and option expected values [5, 48, 49].

In addition to assessing subjective values using choices as standardly done in behavioral economics and non-human animal-based neuroscience, we also assessed subjective option values using explicit ratings [31]. Despite the fact that wealth of evidence in decisionmaking research suggests that subjective values are highly dependent on whether they are inferred from choices or ratings (also referred to the revealed versus state preferences dichotomy [50], post-learning explicit ratings delivered results remarkably consistent with choice-revealed preferences. Indeed, transfer phase choices could almost perfectly be reproduced from explicit ratings, which were, in turn, more consistent with range, rather than divisive normalization process. In addition to provide a welcome test of robustness of our results, the similarity between choice-based and rating-based subjective values, also demonstrates the context-dependent valuation span across procedural as well as declarative representational systems [51, 52].

Beyond the specific question of its functional form, one could ask why option values would be learned and represented in a relative or normalized manner? In other terms, what is the functional reason for context-dependent representations? One partial answer to this question can be tracked to studies showing that context-dependent preferences are ecologically rational (in other words, they convey a statistical or strategical advantage over unbiased representations [53]. On a similar vein, it could be proposed that unbiased value representations are computationally more costly, making relative or context-dependent encoding an efficient solution. Consistent with this idea, a recent study indicates that human participants are capable to adaptatively modulate their value representations from relative to absolute, as they learn that the latter scheme is more advantageous [43]. Another study confirmed that values representations are somehow flexible by showing that shifting the attentional focus from subject emotions to perceive outcome shift the balance from relative to unbiased value encoding [36]. However, neither of these studies manage to report situations in which the representational code was fully context-independent. Taking into account that relative value encoding has been shown in a plethora of species and situations [6–8,10,44,45], it seems also reasonable to suppose that these findings stem from some deep preserved principles of how (value-based) information is encoded and represented in the brain. Accordingly, normalization naturally emerges as a solution to maximize the gain function between underlying stimuli (whose range may vary greatly) and neural response [54]. Crucially, and consistently with our results, while there is ample evidence that the divisive normalization rule provides a good account of information encoding in the perceptual system [55], several primate neurophysiological studies indicate the value-related signals in dopaminergic neurons [56] and the orbitofrontal cortex [12, 13]: hubs of the brain valuation system [57,58]. Similar findings have been replicated in human fMRI [22,27,59]. On the other side, neural evidence of divisive normalization in value-based decision-making is scant in the brain valuation system (but see [6]), although it remains possible that activity in the perceptual and attentional systems (such as the parietal cortex) displays signs of divisive normalization [16].

Multiple elements of our results concordantly show that divisive normalization does not provide a good account of subjective value representation in human reinforcement learning. It is nonetheless still possible that this rule provides a good description of human behavior in other value-based decision-making domains. In fact, previous studies claiming evidence for divisive normalization used other tasks involving items whose values have not to be extracted from experience17,19,60. However, it is worth noting that evidence concerning previous reports of divisive normalization in humans have been recently challenged and alternatively accounts, such as range normalization, have not been properly tested in these datasets23,61. It is worth mentioning that range normalization principle has been recently successfully adapted to account for decision-making under risk62. We therefore believe that our and other recent findings strongly call for radically rethinking the putative role of divisive normalization in value-based decision-making comparing with alternative models of normalization and using highly diagnostic experimental designs, as the one used here. On the other side, by using reinforcement learning framework, our design has the advantage that it can be readily translated into animal research to further extend and characterize the neural mechanisms underlying these findings. Further research will also determine whether range normalization rule also applies to primary (positive and negative) rewards, even if previous evidence (in human and animals) suggest that contextdependent principles apply to food and electric shocks [6–8, 10, 44, 45, 60].

Despite the fact the range normalization presents several computational advantages compared to divisive normalization (such as the fact of being easily translatable to partial feedback and punishment avoidance tasks [4, 26], we believe there is still room for improvement. For instance, the weighted range normalization rule, although descriptively accurate, is silent concerning its cognitive origin mechanism. Future research, for example feature eye-tracking, will be needed to shed light on these aspects. Future research will also be needed to assess to which extent the same rule applies to vast decision spaces involving more than three options.

To conclude, while our results cast serious doubt about the relevance of divisive normalization principle in value-based decisionmaking [33], they also establish once again that context-dependence represents one of the most pervasive feature of human cognition and provide further insights into its algorithmic instantiation.

## Materials and methods

### Participants

We recruited 300 participants (126 females, 148 males, 26 N/A, aged 26.09±7.88 years old) via the Prolific platform (www.prolific.co). The research was carried out following the principles and guidelines for experiments including human participants provided in the declaration of Helsinki (1964, revised in 2013). The INSERM Ethical Review Committee / IRB00003888 approved and participants were provided written informed consent prior to their inclusion. The results presented in the main text are those of Experiment 1 only (N=150). The results of an alternative design (Experiment 2) are presented in the Supplementary Materials. To sustain motivation throughout the experiment, participants were given a bonus depending on the number of points won in the experiment (average money won in pounds: 5.05±0.50, average performance against chance during the learning phase and transfer phase: M=0.74±0.087, t(149)=34.04, p¡0.0001, d=2.78). The data of one participant for the explicit phase was not included due to technical issues. A pilot online-based experiment was originally performed (N=40, the results are also presented in the Supplementary Materials).

### Behavioral task

Participants performed an online version of a probabilistic instrumental learning task, instantiated as a multi-armed bandit task. After checking the consent form, participants received written instructions explaining how the task worked and that their final payoff would be affected by their choices in the task. During the instructions the possible outcomes in points (from 0 to 100 points) were explicitly showed as well as their conversion rate (1 point = 0.02 pence). The instructions were concluded with a short 3-item quiz to make sure participants’ understanding of the task was sufficient. The instructions were then followed by a short training session of 12 trials aiming at familiarizing the participants with the response modalities. If participants’ performance during the training session did not reach 60% of correct answers (i.e., choices towards the option with the highest expected value), they had to repeat the training session. Participants could also voluntarily repeat the training session up to two times and then started the actual experiment.

In our task, options were materialized by abstract stimuli (cues) taken from randomly generated identicons, colored such that the subjective hue and saturation were very similar according to the HSLUV color scheme (www.hsluv.org). On each trial, two or three cues were presented on different positions (left/middle/right) on the screen. The position of a given cue was randomized, such that a given cue was presented an equal number of times on the left, the middle, and the right. Participants were required to select between the cues by clicking on one cue. The choice window was self-paced. A brief delay after the choice was recorded (500ms); the outcome was displayed for 1000ms. There was no fixation screen between trials.

#### Experimental design, version a

The full task consisted in three phases: one learning phase and two elicitation phases. During the learning phase, cues appeared in fixed pairs/triplets. Each pair/triplet was presented 45 times, leading to a total of 180 trials. Within each pair/triplet, the cues were associated to a deterministic outcome drawn from a normal distribution with variable means *µ* ∈ [0, 100] and fixed variance *σ*^2^ = 4 (**Table 1**). At the end of the trial, the cues disappeared and were replaced by the outcome. Once they had completed the learning phase, participants were displayed with the total points earned and their monetary equivalent. After the learning phase, participants performed two elicitation phases: a transfer phase and an explicit rating phase. The order of the elicitation phases was counterbalanced across participants. In the transfer phase, the 10 cues from the learning phase were presented in all possible binary combinations (45, not including pairs formed by the same cue). Each pair of cues was presented four times, leading to a total of 180 trials. Participants were explained that they would be presented with pairs of cues taken from the learning phase, and that all pairs would not have been necessarily displayed together before. On each trial, they had to indicate which of the cues was the one with the highest value. In the explicit rating phase, each cue from the learning phase was presented alone. Participants were asked what was the average value of the cue and had to move a cursor ranging from 0 to 100. Each cue was presented four times, leading to a total of 40 trials. In both elicitation phases, the outcome was not provided in order not to modify the subjective option values learned during the learning phase, but participants were informed that their choices would count for the final payoff.

#### Experimental design, versions b and c

In the learning phase, we added forced-choice trials to the 180 free-choice trials [38]. In these forced trials, only one option was selectable and the other cue(s) were shaded. We added 5 forced-choice trials per option, leading to a total of 230 trials in the learning phase. In version b, even in the forced choice trial, the participants could only see the outcomes of all options. In version c, participants could only see the outcome of the chosen option. The elicitation phases (transfer and explicit rating) remained unchanged.

### Behavioral analyses

For all experiments, we were interested in three different variables reflecting participants’ learning: (1) correct choice rate in the learning phase, i.e., choices directed towards the option with the highest objective value; (2) choice rate in the transfer phase, i.e., the number of times an option is chosen, divided by the number of times the option is presented; (3) subjective valuation in the explicit phase, i.e., average reported value per option. Statistical effects were assessed using multiple-way repeated measures analyses of variance (ANOVAs) with range amplitude (narrow or wide) and number of presented options (binary or trinary decision problem, **Figure 1**) as within-participant factor, and experiment version (a,b,c) as between-participant factors. Post-hoc tests were performed using one-sample and two-sample t tests for respectively within- and between-experiment comparisons. To assess overall performance, additional one-sample t tests were performed against chance level (0.5 – two options contexts – and 0.33 – three options contexts). We report the *t* statistic, *p*-value, and Cohen’s *d* to estimate effect size (two-sample *t*-test only). Given the large sample size (n = 300), central limit theorem allows us to assume normal distribution of our overall performance data and to apply properties of normal distribution in our statistical analyses, as well as sphericity hypotheses. Concerning ANOVA analyses, we report Levene’s test for homogeneity of variance, the uncorrected statistical, as well as Huynh-Feldt correction for repeated measures ANOVA when applicable [61], *F* statistic, *p*-value, partial eta-squared 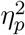, and generalized eta-squared *η*^2^ (when Huynh-Feldt correction is applied) to estimate effect size. All statistical analyses were performed using MATLAB (www.mathworks.com) and R (www.r-project.org).

### Computational models

The goal of our models is to estimate the subjective value of each option, and to choose the option which maximizes the expected reward (in our case, with the highest expected value). At each trial t, the expected value Q of each option is updated with a delta rule:

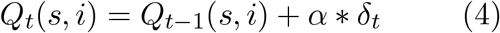

where *α* is the learning rate and *δ*_*t*_ is a prediction error term. At each trial, the chosen and unchosen options are updated with two distinct learning rates and separate, outcome-specific, prediction error terms *δ*_*t*_, calculated as the difference between the subjective outcome *s*(*R*_*i*_) and the expected one:

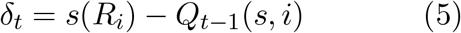

We modeled participants’ choice behavior using a softmax decision rule representing the probability for a participant to choose one option a over the other options one alternative in binary contexts (n=2), two in trinary contexts (n=3):

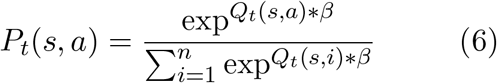

where *n* is the number of outcomes presented in a given trial (*n* ∈ {2; 3}), *β >* 0 is the inverse temperature parameter. High temperatures (*β* → 0) cause the action to be all (nearly) equiprobable. Low temperatures (*β* → +∞) cause a greater difference in selection probability for actions that differ in their value estimates [62].

We compared four alternative computational models: the unbiased (UNBIASED) model, which encodes outcomes on an absolute scale independently of the choice context in which they are presented; the range normalization (RANGE) model, where the reward is normalized as a function of the range of the outcomes, the divisive normalization (DIVISIVE) model, where the reward is normalized as a function of the sum of all the outcomes), and the non-linear range normalization (RANGE^*ω*^) model, where the normalized outcome is power-transformed with an additional free parameter.

### Unbiased model

At trial *t* = 0, for all contexts *Q*_*t*=0_ = 50. For each option *i*, the subjective values *s*(*R*_*i*_) are encoded as the participants see the outcomes, i.e., their objective value in points.

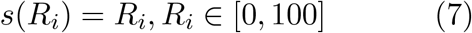

### Range normalization model

At trial *t* = 0, for all contexts *Q*_*t*=0_ = 0.5. The subjective values *s*(*R*_*i*_) are encoded depending on the value of the other options, specifically the maximum and the minimum available rewards.

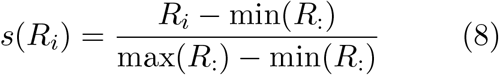

where max(*R*:) and min(*R*:) are, respectively, the maximum and minimum outcome presented in a given trial. In version c, where only the reward of the chosen option is displayed, the outcomes of unchosen options are replaced with the last seen outcomes for these options [5].

### Divisive normalization model

At trial *t* = 0, for all options *Q*_*t*=0_ = 0.5. The outcomes are encoded depending on the value of all the other options, specifically the sum of all available rewards.

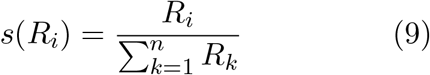

where *n* is the number of outcomes presented in a given trial (*n* ∈ {2; 3}). In version c, where only the reward of the chosen option is displayed, the outcomes of unchosen options are replaced with the last seen outcomes for these options [5].

### Non-linear range normalization model

At trial *t* = 0, for all contexts *Q*_*t*=0_ = 0.5. The subjective values *s*(*R*) are encoded depending on the value of the other options, specifically the maximum and the minimum available rewards. This normalized outcome is then set to the power of *ω*, with 0 *< ω <* +∞:

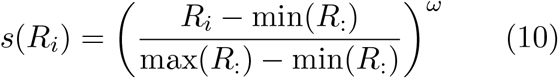

where min(*R*:) and min(*R*:) are, respectively, the maximum and minimum outcome presented in a given trial. In version c, where only the reward of the chosen option is displayed, the outcomes of unchosen options are replaced with the last seen outcomes for these options [5].

### Ex-ante simulations

The model predictions displayed in **Figure 1C** were obtained by simulating choices of artificial agents. The update rule for the option values is described in **Equations 1 and 2**.

Predictions for each experiment were simulated for a set of 50 agents, to match our number of participants per version. Each agent was associated with a set of parameters [*β, α*_*c*_, *α*_*u*_] for the inverse temperature, the learning rate of the chosen option, the learning rate for unchosen options, respectively. The parameters were independently drawn from prior distributions, which we took to be Beta(1.1,1.1) for the learning rates and Gamma(1.2,5) for the inverse temperature [63]. The value of the inverse temperature is irrelevant in the learning phase, because the feedback is always complete, which means that the options should converge to their subjective average value independently of the choice, provided that the learning rates are different from 0. Moreover, in the transfer phase, to obtain the agents’ preferences based on the learned option values, we chose to use an argmax decision rule instead of a softmax decision rule. At each trial t in the transfer phase, comparing option a and option b, the probability of choosing option a is calculated as follows

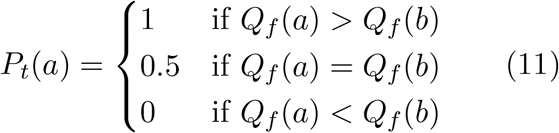

where *Q*_*f*_ is a vector of the final Q-values at the end of the learning phase.

## Acknowledgments

SP is supported by the Institut de Recherche en Santé Publique (IRESP, grant number: 20II138-00), and the Agence National de la Recherche (CogFinAgent: ANR-21-CE230002-02; RELATIVE: ANR-21-CE37-0008-01; RANGE : ANR-21-CE28-0024-01). The Department d’études cognitives is supported by the Agence National de la Recherche (ANR; FrontCog ANR-17-EURE-0017). SB acknowledges support from the European Research Council (ERC) under the European Union’s Horizon 2020 research and innovation program (Grant agreement No. 948545).

## Supplementary Materials

### Pilot experiment with noise-less rewards

To ascertain that our task design would be feasible and that participants would be able to learn the values of ten options by trial-and-error, a pilot online-based experiment was originally performed. We recruited 40 participants (23 females, 16 males, 1 N/A, aged 30.35±9.73 years) via Prolific (www.prolific.co). In the pilot experiments, the outcome variance was set to σ = 0, i.e., the rewards were displayed without any noise (in the main tasks, the variance was set to σ = 4). In order to characterize learning behavior of participants, we analyzed the correct response rate in the learning and the transfer phases, i.e., choices directed toward the most favorable option at each trial. To assess successful learning, we first tested participants’ correct response rate against chance level. We found it to be above chance level in both the learning phase (0.5 and 0.33 in the binary and trinary learning contexts, respectively; Pilot experiment 1: *t*(19)=11.83, *p*<.0001, *d*=2.65, **Figure S1A**; Pilot experiment 2: *t*(19)=7.39, *p*<.0001, *d*=1.65, **Figure S1B**) and the transfer phase (0.5; Pilot experiment 1: *t*(19)=5.87, *p*<.0001, *d*=1.31, **Figure S1A**; Pilot experiment 2: *t*(19)=3.01, *p*=.0072, *d*=0.67, **Figure S1B**).

**Figure S1.**
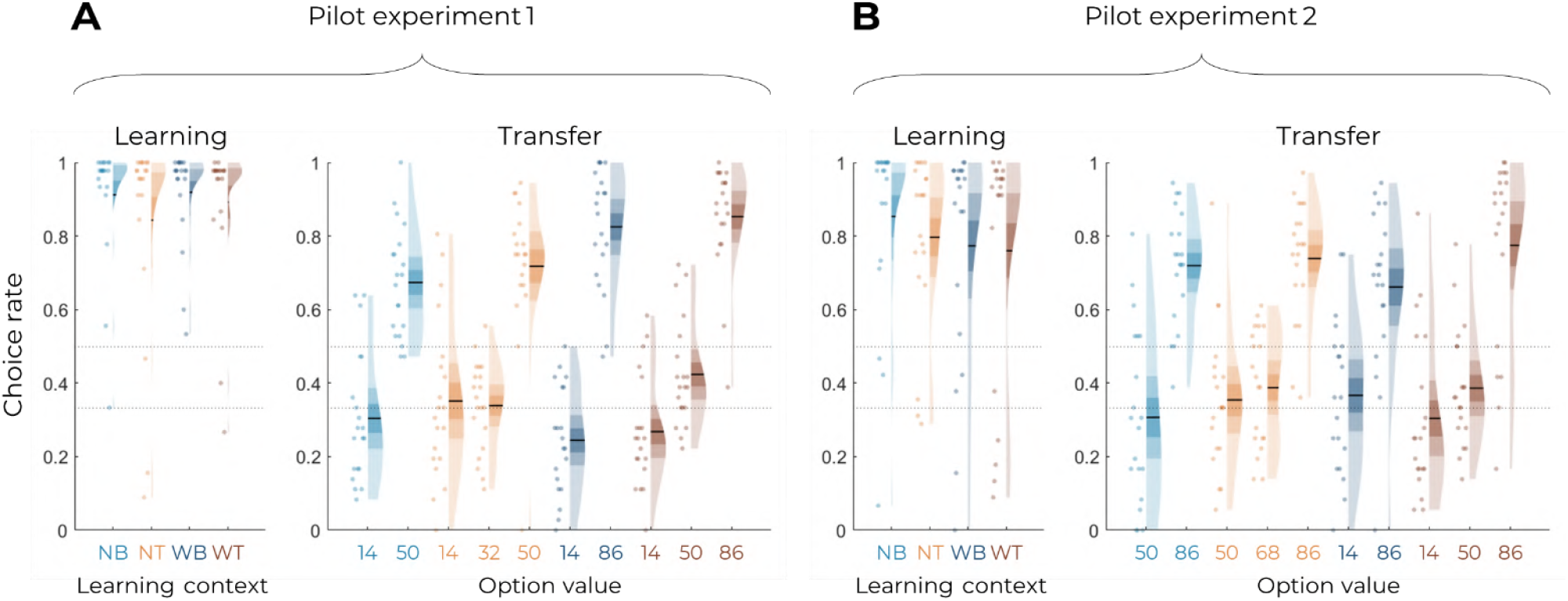
Behavioral results of the pilot experiments. Correct choice rate in the learning phase as a function of the choice context (left panels), and choice rate per option in the transfer phase (right panels) for Pilot Experiment 1 **(A)** and Pilot Experiment 2 **(B)**. In all panels, points indicate individual average, shaded areas indicate probability density function, 95% confidence interval, and SEM. NB: narrow binary, NT: narrow trinary, WB: wide binary, WT: wide trinary.

### Experiment 2

#### Experimental design

In addition to Experiment 1, whose results are presented in the main text, we recruited 150 participants to perform a modified version of Experiment 1. The only difference between Experiment 1 and Experiment 2 is the value of the options in the narrow contexts: in Experiment 1, they went from 14 to 50; in Experiment 2, they went from 50 to 86 (**Figure 1A-B, Figure S2A-B**). Similar to Experiment 1, participants were given a bonus depending on the number of points won in the experiment (average money won in pounds: 6.38±0.58, average performance against chance during the learning phase and transfer phase: M=0.78±0.099, *t*(149)=34.65, *p*<0.0001, *d*=2.83). No data had to be excluded for technical issues.

**Figure S2.**
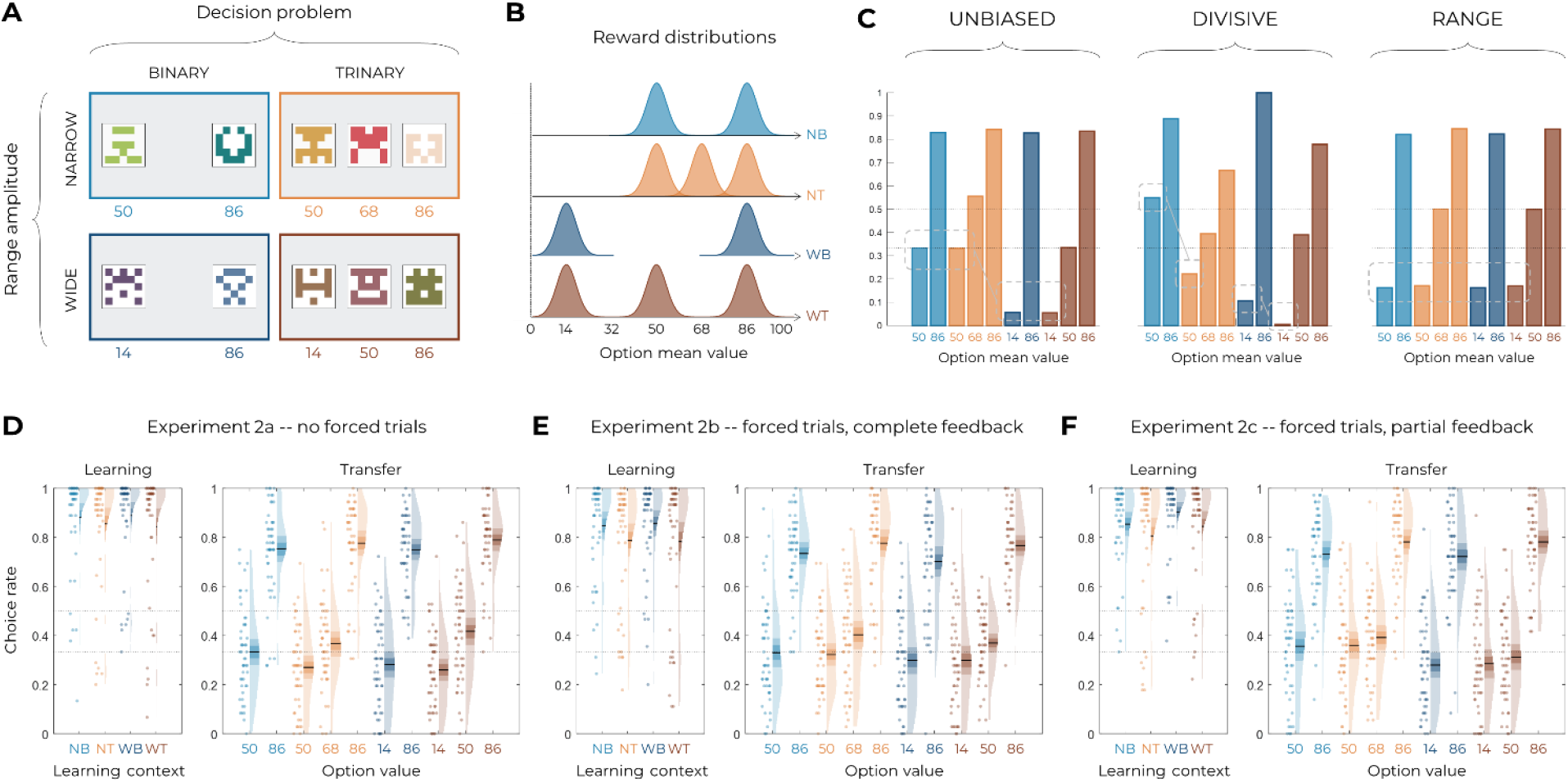
Experimental design, model predictions, and behavioral results. **(A)** Choice contexts in the learning phase. Participants were presented with four choice contexts varying in the amplitude of the outcomes’ range (narrow or wide) and the number of options (binary or trinary decisions). **(B)** Means of each reward distribution. After each decision, the outcome of each option was displayed on the screen. Each outcome was drawn from a normal distribution with variance σ^2^ = 4. NB: narrow binary, NT: narrow trinary, WB: wide binary, WT: wide trinary. **(C)** Model predictions of the transfer phase choice rates for the UNBIASED (left), DIVISIVE (middle) and RANGE (right) models. Dashed lines represent the key prediction for each model. **(D-F)** Correct choice rate in the learning phase as a function of the choice context (left panels), and choice rate per option in the transfer phase (right panels) for the three versions of the main experiment: without forced trials **(D)**, with forced trials and complete feedback information **(E)** and with forced trials and partial feedback information **(F)**. In all panels, points indicate individual average, shaded areas indicate probability density function, 95% confidence interval, and SEM. NB: narrow binary, NT: narrow trinary, WB: wide binary, WT: wide trinary.

#### Behavioral results

In the learning phase, the correct response rate was significantly higher than chance level (0.5 and 0.33 in the binary and trinary learning contexts, respectively) in all conditions (least significant: *t*(49)=13.71, *p*<.0001, *d*=1.94; on average: t(49)=15.75, p<.0001, d=2.23; **Figure S2D**). We further checked whether the task factors affected performance in the learning phase and found a small significant effect of the decision problem (the correct choice rate being higher in the binary compared to the trinary contexts: *F*(1,49)=4.11, *p*=.048, η^2^ =0.08), but no effect of range amplitude (wide versus narrow; *F*(1,49)=0.027, *p*=.87, η^2^ =0.00) or interaction (*F*(1,49)=0.92, *p*=.34, η^2^ =0.02). In the transfer phase, the correct choice rate in the transfer was significantly higher than chance (*t*(49)=9.56, *p*<.0001, *d*=1.35), thus providing positive evidence of value retrieval and generalization^1–3^. Contrary to what was predicted by the UNBIASED or the DIVISIVE models, the choice rate for the lowest value options (NB_50_, NT_50_, WB_14_ and WT_14_) did not follow the patterns depicted in **Figure S2C**. In fact, all lowest value options displayed a similar choice rate (*F*(3,49)=1.85, *p*=.14, η^2^ =0.04), which is only consistent with the predictions of the RANGE model. Concerning other features of the transfer phase performance, the mid value options valuation is also consistent with the RANGE model, which predict that these options will be valued equally (NT_68_ and WT_50_; *t*(49)=1.34, *p*=.19, *d*=-0.19), contrary to the UNIBIASED and DIVISIVE models which both predict NT_68_ to be greater than WT_50_. Similar to Experiment 1, these mid value options displayed a choice rate very close to that of the corresponding lowest value options (NT_50_ and WT_14_): this feature is still not perfectly captured by the RANGE model (which predicts their choice rate perfectly in between those of high and low value options). To rule out that this effect was not due to a lack of attention for the low and mid value options, we also designed two additional experiments where we added forced choice trials to focus the participants’ attention on all possible options (**Table 1**). As in Experiment 1, focusing participants’ attention to all possible outcomes by forcing their choice did not significantly affect the behavioral performance neither in the learning phase (*F*(2,147)=0.78, *p*=.46, η^2^ =0.01, Levene’s test *F*(2,147)=0.38, *p*=.69) nor in the transfer phase (*F*(2,147)=0.81, *p*=.45, η^2^ =0.01, Levene’s test *F*(2,147)=0.81, *p*=.45). Given the absence of detectable differences across experiments, in the model-based analyses that follow, we pooled the three experiments together. To sum up, the behavioral results are consistent with those of Experiment 1, are in contrast with the predictions of both the UNBIASED and the DIVISIVE models and are rather consistent with the range normalization process proposed by the RANGE model.

#### Model comparison and model simulations

Similar to Experiment 1, model comparison favored the RANGE model over to both the DIVISIVE and the UNBIASED models (oosLL_RAN_ vs oosLL_DIV_: *t*(149)=10.61, *p*<.0001, *d*=0.87; oosLL_RAN_ vs oosLL_UNB_: *t*(149)=5.71, *p*<.0001, *d*=0.47; **Table 2**). Model simulations also confirmed what inferred from the *ex-ante* simulations and indicated that the RANGE model predicts results much closer to the observed ones in respect of many key comparisons (**Figure S3A-C**).

**Figure S3.**
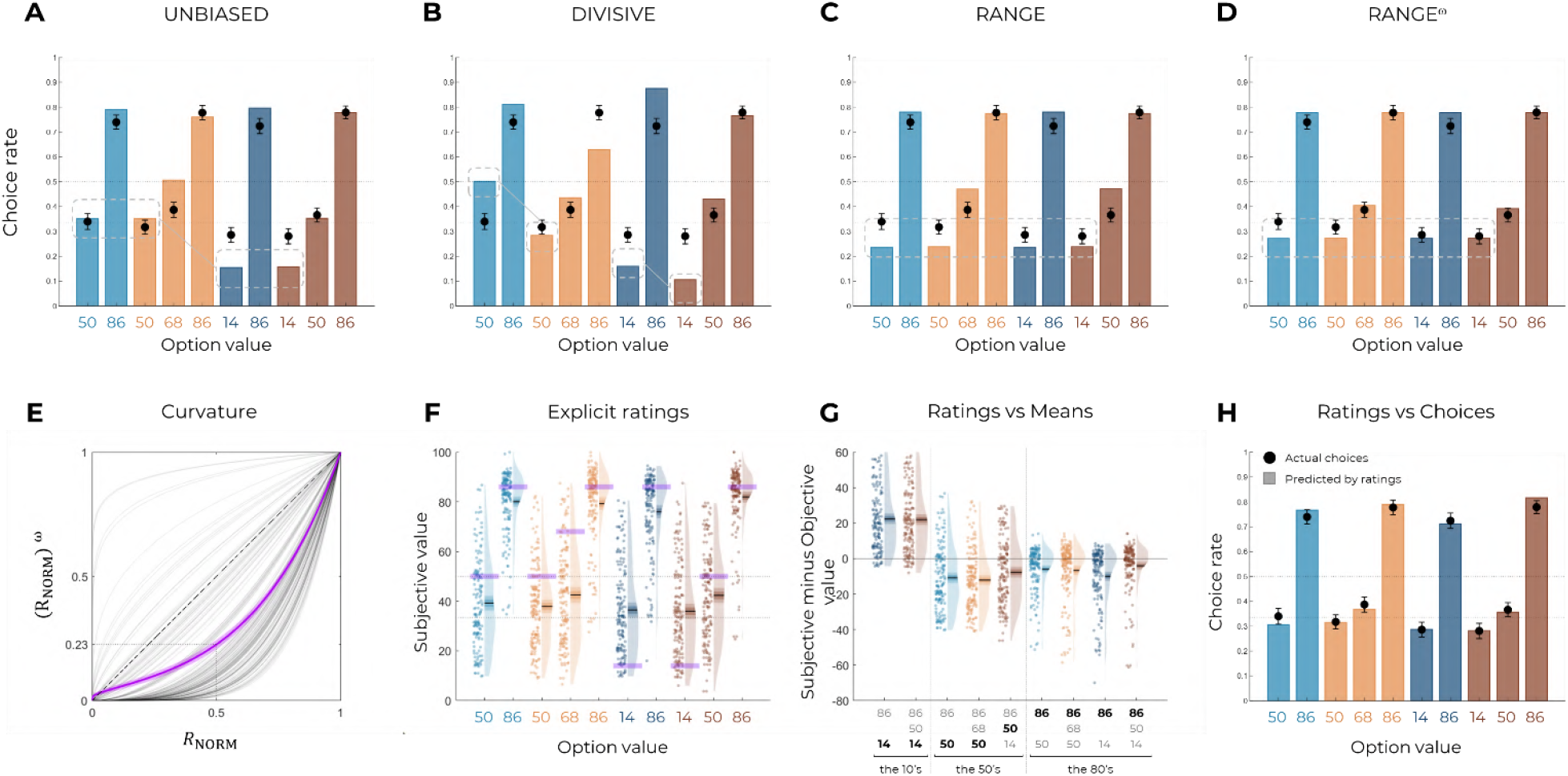
Qualitative model comparison and explicit ratings. Behavioral data (black dots) superimposed on simulated data (colored bars) for the UNBIASED **(A)**, DIVISIVE **(B)**, RANGE **(C)** and RANGE^ω^ **(D)** models. Simulated data in the transfer phase were obtained with the bestfitting parameters, optimized on all four contexts of the learning phase. Dashed lines represent the key prediction for each model. **(E)** Curvature function of the normalized reward per participant. Each grey line was simulated with the best-fitting power parameter ω for each participant. Dashed line represents the identity function (ω = 1), purple line represents the average curvature over participant, shaded area represents SEM. **(F)** Reported subjective values in the elicitation phase for each option. Points indicate individual average, shaded areas indicate probability density function, 95% confidence interval, and SEM. Purple areas indicate the actual objective value for each option. **(G)** Difference between reported subjective value and actual objective value for each option, arrange in ascending order. The legend of the x-axis represents the values of the options of the context in which each option was learned (actual option value shown in bold black). Points indicate individual average, shaded areas indicate probability density function, 95% confidence interval, and SEM. **(H)** Behavioral choice-based data (black dots) superimposed on simulated choice-based data (colored bars). Simulated data were obtained with an argmax rule assuming that subjects were making transfer phase decisions based on the explicit subjective ratings of each option.

#### Improving the RANGE model

Model comparison and model simulations of the RANGE^ω^ model supported the conclusions from Experiment 1. On average, the power parameter ω was greater than 1 (mean±std: 2.83±1.47, *t*(149)=15.25, *p*<.0001, *d*=1.25), suggesting that participants value the mid value options less than the midpoint between lowest and highest options (i.e., closer to the lowest option, **Figure S3E**). Quantitative model comparison favored the RANGE^ω^ model over all other models, including the RANGE model (**Table 2**) (oosLL_RAN_ vs oosLL_RAN(ω)_ : *t*(149)=-8.63, *p*<.0001, *d*=-0.70; **Table 2**). Moreover, the inspection of model simulations confirmed that the RANGE^ω^ model perfectly captures participants’ behavior in the transfer phase. More specifically, the mid value options (NT_68_ and WT_50_) and the lowest value options (NB_14_, NT_14_, WB_14_ and WT_14_) were better estimated in all contexts (**Figure S3D**). To conclude, the addition of a power parameter allowed our model to match participants’ behavior almost perfectly.

#### Explicit assessment of option values

Consistently with the results of Experiment 1, the subjective values elicited through explicit ratings were consistent with those elicited though binary choices in many key aspects (**Figure S3F-G**). Again, to compare elicitation methods, we simulated transfer phase choices based on the explicit elicitation ratings. We found the pattern simulated using explicit ratings to perfectly match the actual choice rates of the transfer phase (*t*(149)=1.18, *p*=.24, *d*=0.10, **Figure S2H**), suggesting that both elicitation methods tap into the same valuation system.

### Ruling out alternative formulations of the divisive normalization

#### DIVISIVE^ω^ model

To confirm that our power manipulation in the RANGE^ω^ model would not affect our predictions in the DIVISIVE model (especially, the difference in value between options from binary and trinary contexts, e.g., WB_86_ and WT_86_), we implemented the same manipulation in the DIVISIVE model. In the DIVISIVE^ω^ model, the normalized outcome is power-transformed by the ω parameter (0 < ω < +∞), as follows:

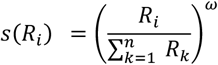

where *n* is the number of contextually relevant stimuli. Crucially, for ω = 1, the DIVISIVE^ω^ model reduces to the DIVISIVE model. As expected, the DIVISIVE^ω^ model was unable to match participants’ behavior, despite a small improvement in the prediction of the choice rates for mid and lowest value options, in both experiments (**Figure S4B,E**). Moreover, over both experiments, quantitative model comparison favored the DIVISIVE^ω^ model over the DIVISIVE model (oosLL_DIV(ω)_=-129.25±64.94, median=-116.98; oosLL_DIV(ω)_ vs oosLL_DIV_: *t*(299)=11.50, *p*<.0001, *d*=0.66) but not over range-adaptation model (oosLL_DIV(ω)_ vs oosLL_RAN(ω)_: *t*(299)=-11.71, *p*<.0001, *d*=-0.68). In conclusion, the addition of a power transformation was insufficient to correct the behavioral predictions of the DIVISIVE model.

**Figure S4.**
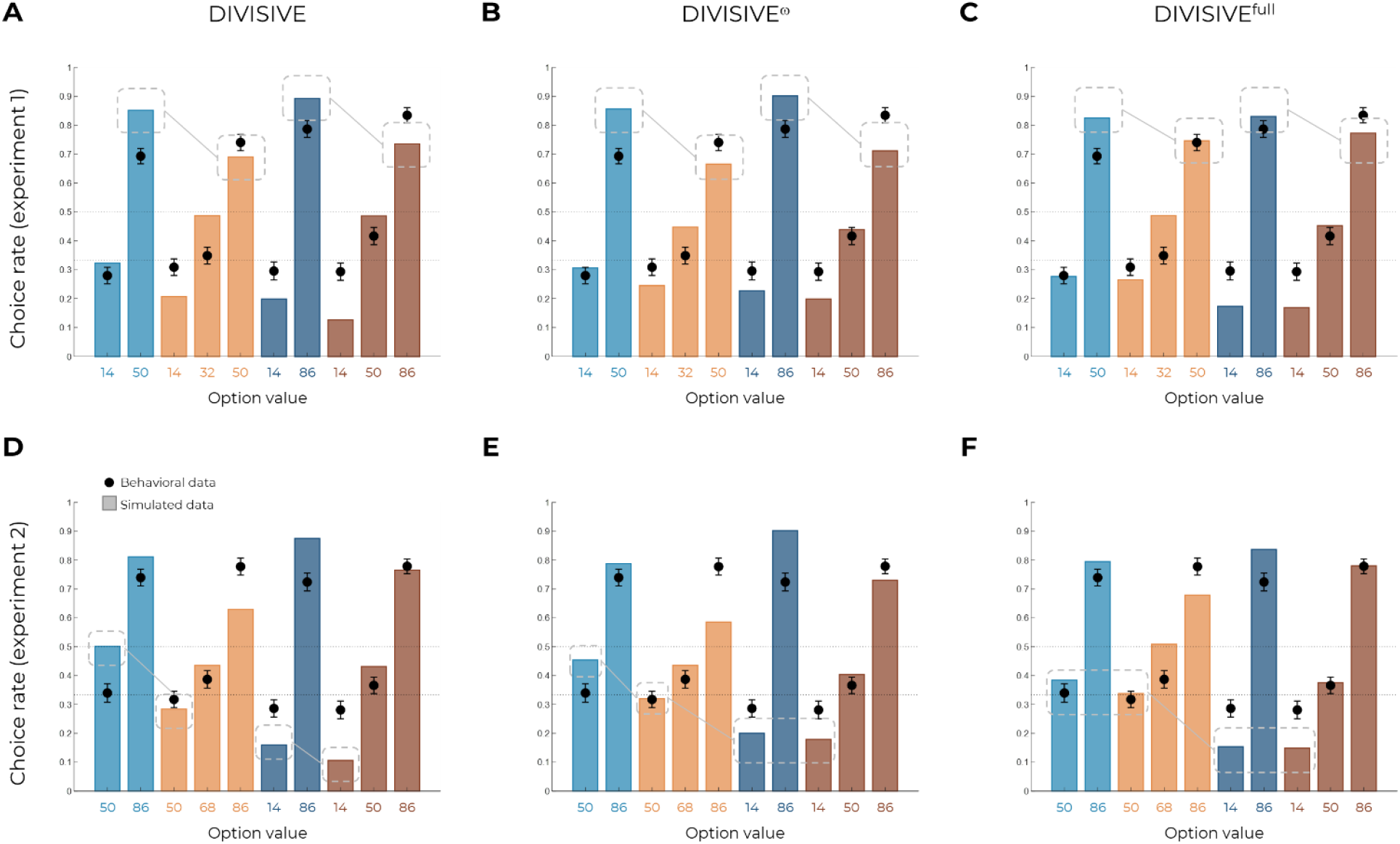
Ruling out divisive normalization. Behavioral data (black dots) superimposed on simulated data (colored bars) for Experiment 1 (top row) and Experiment 2 (bottom row), for the DIVISIVE **(A, D)**, DIVISIVE^ω^ **(B, E)** and DIVISIVE^full^ **(C, F)** models. Simulated data in the transfer phase were obtained with the best-fitting parameters, optimized on all four contexts of the learning phase. Dashed lines represent key features that allow discriminating divisive normalization models from range normalization and unbiased value representations.

#### DIVISIVE^full^ model

Finally, we acknowledge that the normalization rule we implemented is a simpler implementation of classical divisive normalization^4–6^. To make sure that this over-simplification did not affect the main results of this study, we implemented a more complex version of divisive normalization, including a semi-saturation parameter, a normalization weight parameter, and a *p*-norm parameter^6^:

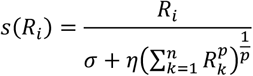

where *n* is the number of contextually relevant stimuli; the parameter 0 < σ < +∞ determines, in a neural system, how neural activity saturates with increased input and can be interpreted as the baseline activity level in the normalization; the parameter 0 < *η* < 1 determines the contribution to the normalization from other alternatives; each alternative is scaled by the magnitude and number of its elements by a norm of degree 1 < *p* < +∞. Crucially, the DIVISIVE^full^ model is nested within the DIVISIVE and UNBIASED model: when σ = 0, *η* = 1 and *p* = 1, the DIVISIVE^full^ model reduces to the DIVISIVE model; when σ = 1 and *η* = 0, the DIVISIVE^full^ model reduces to the UNBIASED model (no normalization). Quantitative model comparison showed a substantial improvement in the ability of the DIVISIVE^full^ model to fit participants’ data over the simple DIVISIVE model (oosLL_DIV(full)_=-136.19±88.33, median=-120.63; oosLL_DIV(full)_ vs oosLL_DIV_: *t*(299)=4.69, *p*<.0001, *d*=0.27), but not over range adaptation models (oosLL_DIV(full)_ vs oosLL_RAN(ω)_: *t*(299)=-10.36, *p*<.0001, *d*=-0.60). These comparisons were consistent with model simulations, which clearly show that the key features of the model fail to predict transfer performance (**Figure S4C,F**). In conclusion, and unsurprisingly given the structure of the model, the more complex version of the divisive normalization rule was unable to match participants’ behavior in our task.

**Figure S5.**
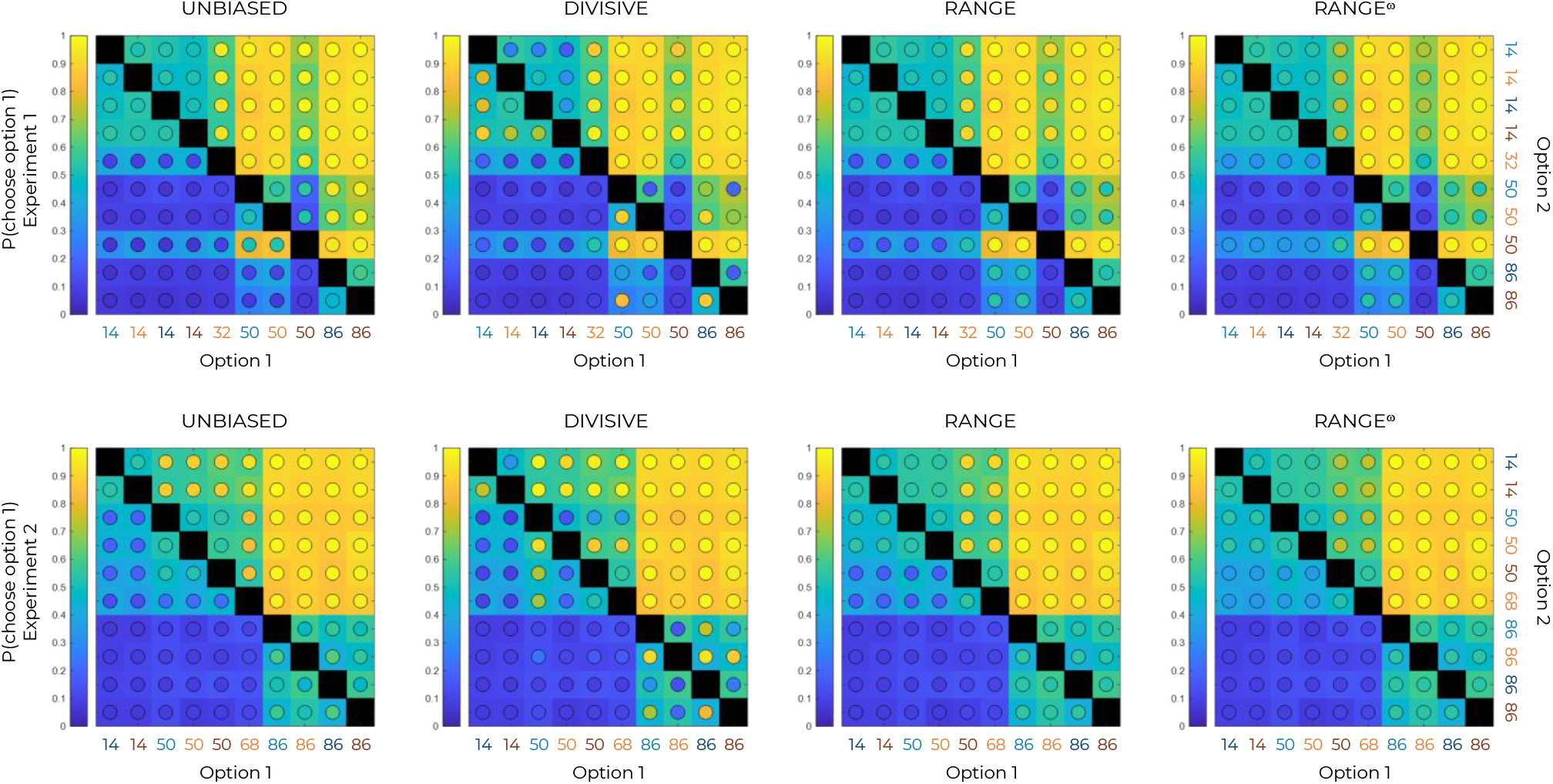
Choice rates per option in the transfer phase and model simulations. Colored maps of pairwise choice rates during the transfer phase of Experiment 1 (top row) and experiment 2 (bottom row), for each option when compared to each of the nine other options, noted here generically as Option 1 and Option 2 in increasing order. Model simulations (colored circles) are superimposed over behavioral data (colored squares). Comparisons between the same symbols are undefined (black squares).

## Footnotes

1 As clearly put but the main advocate of the divisive normalization rule for value-based decion-making: “[…] a system would be highly sensitive to the number of options under consideration. As the number of elements in the denominator grows, so does the aggregate value of the denominator, shifting the overall firing rates lower and lower” [33] (Paul Glimcher, TICS, 2022; page 14)

## Notes

### Competing Interest Statement

The authors have declared no competing interest.

### Summary of Updates

Affiliations corrected

## References

[1] Daniel Kahneman and Amos Tversky. Choices, Values, and Frames. American Psychologist, (39):341–350, 1984.

[2] Joel Huber, John W. Payne, and Christopher Puto. Adding Asymmetrically Dominated Alternatives: Violations of Regularity and the Similarity Hypothesis. Journal of Consumer Research, 9(1):90–98, 1982. Publisher: Oxford University Press.

[3] Tilmann A. Klein, Markus Ullsperger, and Gerhard Jocham. Learning relative values in the striatum induces violations of normative decision making. Nature Communications, 8:16033, June 2017.

[4] Sophie Bavard, Maël Lebreton, Mehdi Khamassi, Giorgio Coricelli, and Stefano Palminteri. Reference-point centering and range-adaptation enhance human reinforcement learning at the cost of irrational preferences. Nature Communications, 9(1):4503, 2018.

[5] Mikhail S. Spektor, Sebastian Gluth, Laura Fontanesi, and Jörg Rieskamp. How similarity between choice options affects decisions from experience: The accentuation-of-differences model. Psychological Review, 126(1):52–88, January 2019.

[6] Hiroshi Yamada, Kenway Louie, Agnieszka Tymula, and Paul W. Glimcher. Free choice shapes normalized value signals in medial orbitofrontal cortex. Nature Communications, 9(1):162, January 2018.

[7] Katherine E. Conen and Camillo Padoa-Schioppa. Partial Adaptation to the Value Range in the Macaque Orbitofrontal Cortex. Journal of Neuroscience, 39(18):3498–3513, May 2019.

[8] Lorena Pompilio and Alex Kacelnik. Context-dependent utility overrides absolute memory as a determinant of choice. Proceedings of the National Academy of Sciences of the United States of America, 107(1):508–512, January 2010.

[9] Lorena Pompilio, Alex Kacelnik, and Spencer T. Behmer. State-dependent learned valuation drives choice in an invertebrate. Science (New York, N.Y.), 311(5767):1613–1615, March 2006.

[10] Cwyn Solvi, Yonghe Zhou, Mark Roper, Yunxiao Feng, Li Sun, Rebecca Reid, Lars Chittka, Andrew Barron, and Fei Peng. Bumblebees retrieve only the ordinal ranking of foraging options when comparing memories obtained in distinct settings, April 2022. Pages: 2022.04.05.487177 Section: New Results.

[11] A. L. Fairhall, G. D. Lewen, W. Bialek, and R. R. de Ruyter Van Steveninck. Efficiency and ambiguity in an adaptive neural code. Nature, 412(6849):787–792, August 2001.

[12] Camillo Padoa-Schioppa. Range-adapting representation of economic value in the orbitofrontal cortex. The Journal of Neuroscience: The Official Journal of the Society for Neuroscience, 29(44):14004–14014, November 2009.

[13] Shunsuke Kobayashi, Ofelia Pinto de Carvalho, and Wolfram Schultz. Adaptation of Reward Sensitivity in Orbitofrontal Neurons. The Journal of Neuroscience, 30(2):534–544, January 2010.

[14] Kenway Louie and Paul W. Glimcher. Efficient coding and the neural representation of value. Annals of the New York Academy of Sciences, 1251:13–32, March 2012.

[15] John H. Reynolds and David J. Heeger. The normalization model of attention. Neuron, 61(2):168– 185, January 2009.

[16] Kenway Louie, Lauren E. Grattan, and Paul W. Glimcher. Reward Value-Based Gain Control: Divisive Normalization in Parietal Cortex. Journal of Neuroscience, 31(29):10627–10639, July 2011. Publisher: Society for Neuroscience Section: Articles.

[17] Kenway Louie, Mel W. Khaw, and Paul W. Glimcher. Normalization is a general neural mechanism for context-dependent decision making. Proceedings of the National Academy of Sciences of the United States of America, 110(15):6139–6144, April 2013.

[18] Kenway Louie, Paul W Glimcher, and Ryan Webb. Adaptive neural coding: from biological to behavioral decision-making. Current Opinion in Behavioral Sciences, 5:91–99, October 2015.

[19] Ryan Webb, Paul W. Glimcher, and Kenway Louie. The Normalization of Consumer Valuations: Context-Dependent Preferences From Neurobiological Constraints. Management Science, May 2020.

[20] R. J. Herrnstein. Relative and Absolute Strength of Response as a Function of Frequency of Reinforcement1,2. Journal of the Experimental Analysis of Behavior, 4(3):267–272, 1961. eprint:https://onlinelibrary.wiley.com/doi/pdf/10.1901/jeab.1961.4-267

[21] Camillo Padoa-Schioppa and Aldo Rustichini. Rational Attention and Adaptive Coding: A Puzzle and a Solution. American Economic Review, 104(5):507–513, May 2014.

[22] Christopher J. Burke, Michelle Baddeley, Philippe N. Tobler, and Wolfram Schultz. Partial Adaptation of Obtained and Observed Value Signals Preserves Information about Gains and Losses. Journal of Neuroscience, 36(39):10016–10025, September 2016.

[23] Sebastian Gluth, Nadja Kern, Maria Kortmann, and Cécile L. Vitali. Value-based attention but not divisive normalization influences decisions with multiple alternatives. Nature Human Behaviour, 4(6):634–645, June 2020.

[24] Allen Parducci. Range-frequency compromise in judgment. Psychological Monographs: General and Applied, 77(2):1–50, 1963. Place: US Publisher: American Psychological Association.

[25] Aldo Rustichini, Katherine E. Conen, Xinying Cai, and Camillo Padoa-Schioppa. Optimal coding and neuronal adaptation in economic decisions. Nature Communications, 8(1):1208, October 2017.

[26] Sophie Bavard, Aldo Rustichini, and Stefano Palminteri. Two sides of the same coin: Beneficial and detrimental consequences of range adaptation in human reinforcement learning. Science Advances, 7(14):eabe0340, April 2021. Publisher: American Association for the Advancement of Science Section: Research Article.

[27] Karin M. Cox and Joseph W. Kable. BOLD Subjective Value Signals Exhibit Robust Range Adaptation. Journal of Neuroscience, 34(49):16533–16543, December 2014.

[28] Maë l Lebreton, Sophie Bavard, Jean Daunizeau, and Stefano Palminteri. Assessing interindividual differences with task-related functional neuroimaging. Nature Human Behaviour, 3(9):897–905, September 2019.

[29] Seth Roberts and Harold Pashler. How persuasive is a good fit? A comment on theory testing. Psychological Review, 107(2):358–367, 2000. Place: US Publisher: American Psychological Association.

[30] Stefano Palminteri, Valentin Wyart, and Etienne Koechlin. The Importance of Falsification in Computational Cognitive Modeling. Trends in Cognitive Sciences, 21(6):425–433, June 2017.

[31] Basile Garcia, Fabien Cerrotti, and Stefano Palminteri. The description–experience gap: a llenge for the neuroeconomics of decisionmaking under uncertainty. Philosophical Transactions of the Royal Society B: Biological Sciences, 376(1819):20190665, March 2021. Publisher: Royal Society.

[32] Stefano Palminteri, Mehdi Khamassi, Mateus Joffily, and Giorgio Coricelli. Contextual modulation of value signals in reward and punishment learning. Nature Communications, 6:8096, August 2015.

[33] Paul W. Glimcher. Efficiently irrational: deciphering the riddle of human choice. Trends in Cognitive Sciences, 0(0), May 2022. Publisher: Elsevier.

[34] Ralph Hertwig and Ido Erev. The description-experience gap in risky choice. Trends in Cognitive Sciences, 13(12):517–523, December 2009.

[35] Jian Li and Nathaniel D. Daw. Signals in human striatum are appropriate for policy update rather than value prediction. The Journal of neuroscience : the official journal of the Society for Neuroscience, 31(14):5504–5511, April 2011.

[36] William Hayes and Douglas Wedell. Reinforcement-Learning In and Out of Context: The Effects of Attentional Focus. Journal of Experimental Psychology Learning Memory and Cognition, March 2022.

[37] Andrei R. Teodorescu and Marius Usher. Disentangling decision models: From independence to competition. Psychological Review, 120(1):1–38, 2013. Place: US Publisher: American Psychological Association.

[38] Valérian Chambon, Héloïse Théro, Marie Vidal, Henri Vandendriessche, Patrick Haggard, and Stefano Palminteri. Information about action outcomes differentially affects learning from selfdetermined versus imposed choices. Nature Human Behaviour, 4(10):1067–1079, October 2020.

[39] Robert C Wilson and Anne GE Collins. Ten simple rules for the computational modeling of behavioral data. eLife, 8:e49547, November 2019. Publisher: eLife Sciences Publications, Ltd.

[40] Daniel Bernoulli. Specimen Theoriae Novae de Mensura Sortis. 1738.

[41] Elliot A. Ludvig, Christopher R. Madan, Neil McMillan, Yaqian Xu, and Marcia L. Spetch. Living near the edge: How extreme outcomes and their neighbors drive risky choice. Journal of Experimental Psychology: General, 147(12):1905–1918, 2018. Place: US Publisher: American Psychological Association.

[42] Stefano Palminteri and Maël Lebreton. Context-dependent outcome encoding in human reinforcement learning. Current Opinion in Behavioral Sciences, 41:144–151, October 2021.

[43] Keno Juechems, Tugba Altun, Rita Hira, and Andreas Jarvstad. Human value learning and representation reflect rational adaptation to task demands. Nature Human Behaviour, May 2022.

[44] Yukihisa Matsumoto and Makoto Mizunami. Context-Dependent Olfactory Learning in an Insect. Learning & Memory, 11(3):288–293, May 2004. Company: Cold Spring Harbor Laboratory Press Distributor: Cold Spring Harbor Laboratory Press Institution: Cold Spring Harbor Laboratory Press Label: Cold Spring Harbor Laboratory Press Publisher: Cold Spring Harbor Lab.

[45] Lorenzo Ferrucci, Simon Nougaret, Emiliano Brunamonti, and Aldo Genovesio. Effects of reward size and context on learning in macaque monkeys. Behavioural Brain Research, 372:111983, October 2019.

[46] Allen Parducci. Happiness, pleasure, and judgment: The contextual theory and its applications. Happiness, pleasure, and judgment: The contextual theory and its applications. Lawrence Erlbaum Associates, Inc, Hillsdale, NJ, US, 1995. Pages: ix, 225.

[47] Peter P. Wakker. Prospect Theory: For Risk and Ambiguity. Cambridge University Press, 2010.

[48] Ian Krajbich, Carrie Armel, and Antonio Rangel. Visual fixations and the computation and comparison of value in simple choice. Nature neuroscience, 13(10):1292–1298, 2010. Publisher: Nature Publishing Group.

[49] Veronika Zilker and Thorsten Pachur. Nonlinear probability weighting can reflect attentional biases in sequential sampling. Psychological Review, 2021. Publisher: American Psychological Association.

[50] Paul Slovic. The Construction of Preference. Cambridge University Press, Cambridge, 2006.

[51] Samuel J. Gershman and Nathaniel D. Daw. Reinforcement learning and episodic memory in humans and animals: an integrative framework. Annual review of psychology, 68:101–128, January 2017.

[52] Natalie Biderman and Daphna Shohamy. Memory and decision making interact to shape the value of unchosen options. Nature Communications, 12(1):4648, July 2021. Number: 1 Publisher: Nature Publishing Group.

[53] J. M. McNamara, P. C. Trimmer, and A. I. Houston. The ecological rationality of state-dependent valuation. Psychological Review, 119(1):114–119, January 2012.

[54] Horace Barlow. Possible Principles Underlying the Transformations of Sensory Messages. Sensory Communication, 1, January 1961.

[55] Matteo Carandini and David J. Heeger. Normalization as a canonical neural computation. Nature Reviews. Neuroscience, 13(1):51–62, November 2011.

[56] Philippe N. Tobler, Christopher D. Fiorillo, and Wolfram Schultz. Adaptive coding of reward value by dopamine neurons. Science (New York, N.Y.), 307(5715):1642–1645, March 2005.

[57] Oscar Bartra, Joseph T. McGuire, and Joseph W. Kable. The valuation system: A coordinate-based meta-analysis of BOLD fMRI experiments examining neural correlates of subjective value. NeuroImage, 76:412–427, August 2013.

[58] Mathias Pessiglione and Mauricio R. Delgado. The good, the bad and the brain: Neural correlates of appetitive and aversive values underlying decision making. Current Opinion in Behavioral Sciences, 5:78–84, October 2015.

[59] Doris Pischedda, Stefano Palminteri, and Giorgio Coricelli. The Effect of Counterfactual Information on Outcome Value Coding in Medial Prefrontal and Cingulate Cortex: From an Absolute to a Relative Neural Code. Journal of Neuroscience, 40(16):3268–3277, April 2020. Publisher: Society for Neuroscience Section: Research Articles.

[60] Ivo Vlaev, Ben Seymour, Raymond J. Dolan, and Nick Chater. The Price of Pain and the Value of Suffering. Psychological Science, 20(3):309–317, March 2009. Publisher: SAGE Publications Inc.

[61] Ellen R. Girden. ANOVA: Repeated Measures. SAGE, 1992. Google-Books-ID: JomGKpjnfPcC.

[62] Richard S. Sutton and Andrew G. Barto. Reinforcement Learning - An Introduction. Mit Press, 1998.

[63] Nathaniel D. Daw, Samuel J. Gershman, Ben Seymour, Peter Dayan, and Raymond J. Dolan. Model-based influences on humans’ choices and striatal prediction errors. Neuron, 69(6):1204– 1215, March 2011.

## Supplementary references

Bavard, S., Lebreton, M., Khamassi, M., Coricelli, G. & Palminteri, S. Reference-point centering and range-adaptation enhance human reinforcement learning at the cost of irrational preferences. Nat. Commun. 9, 4503 (2018).

Bavard, S., Rustichini, A. & Palminteri, S. Two sides of the same coin: Beneficial and detrimental consequences of range adaptation in human reinforcement learning. Sci. Adv. 7, eabe0340 (2021).

Hayes, W. & Wedell, D. Reinforcement-Learning In and Out of Context: The Effects of Attentional Focus. J. Exp. Psychol. Learn. Mem. Cogn. (2022) doi:10.1037/xlm0001145.

Louie, K., Khaw, M. W. & Glimcher, P. W. Normalization is a general neural mechanism for context-dependent decision making. Proc. Natl. Acad. Sci. U. S. A. 110, 6139–6144 (2013).

Louie, K., Glimcher, P. W. & Webb, R. Adaptive neural coding: from biological to behavioral decision-making. Curr. Opin. Behav. Sci. 5, 91–99 (2015).

Webb, R., Glimcher, P. W. & Louie, K. The Normalization of Consumer Valuations: Context-Dependent Preferences From Neurobiological Constraints. Manag. Sci. (2020) doi:10.1287/mnsc.2019.3536.

